# Systemic Metabolic Rewiring in a Mouse Model of Left Ventricular Hypertrophy

**DOI:** 10.1101/2025.08.13.670184

**Authors:** Alexandra V. Schmidt, Tharika Thambidurai, Olivia D’annibale, Sivakama S. Bharathi, Tim Wood, Eric S. Goetzman, Julian E. Stelzer

## Abstract

Left ventricular hypertrophy (LVH) refers to the pathological thickening of the myocardial wall, and is strongly associated with several adverse cardiac outcomes and sudden cardiac death. While the biomechanical drivers of LVH are well established, growing evidence points to a critical role for cardiac and systemic metabolism in modulating hypertrophic remodeling and disease pathogenesis. Despite the efficiency of fatty acid oxidation (FAO), LVH hearts preferentially increase glucose uptake and catabolism to drive glycolysis and oxidative phosphorylation (OXPHOS). Development of therapies to increase and enhance LFCA FAO are underway, with promising results. However, the mechanisms of systemic metabolic states and LCFA dynamics in the context of cardiac hypertrophy remain incompletely understood. Further, it is unknown to what extent cardiac metabolism is influenced by whole-body energy balance and lipid profiles, despite the common occurrence of lipotoxicity in LVH. In this study, we measured whole-body and cellular respiration along with analysis of lipid and glycogen stores in a mouse model of LVH. We found that loss of the cardiac-specific gene, *Myosin binding protein-C3* (*Mybpc3*), resulted in depletion of adipose tissue, decreased mitochondrial function in skeletal muscle, increased lipid accumulation in both heart and liver, and loss of whole-body metabolic flux. We found that supplementation of exogenous LCFAs boosted LVH mitochondrial function and reversed cardiac lipid accumulation, but did not fully reverse the hypertrophied heart nor systemic metabolic phenotypes. This study indicates that the LVH phenotype caused systemic metabolic rewiring in *Mybpc3^-/-^* mice, and that exogenous LCFA supplementation boosted mitochondrial function in both cardiac and skeletal muscle.

## Introduction

Left ventricular hypertrophy (LVH) is a structural and functional adaptation of the heart in response to increased hemodynamic pressure or volumetric stress caused by increased pressure overload, afterload, or inherited cardiomyopathies. Myocardial wall thickening counteracts increased filling pressures, but oftentimes results in maladaptive remodeling that affects systolic and diastolic function [1]. LVH is commonly associated with comorbidities such as hypertension, obesity, chronic kidney disease, and diabetes [2,3], and although is initially compensatory, sustained LVH results in elevated risks of several adverse cardiac outcomes and sudden cardiac death [2]. While the biomechanical drivers of LVH are well established, growing evidence points to a critical role for cardiac and systemic metabolism in modulating hypertrophic remodeling and disease pathogenesis [4].

Long-chain fatty acids (LCFAs) represent the primary energy substrate for the adult heart under normal physiological conditions, despite the heart’s ability to catabolize many different substrates to fuel oxidative phosphorylation (OXPHOS) in mitochondria. However, during pathological hypertrophy, cardiac substrate utilization shifts away from fatty acid oxidation (FAO)-derived OXPHOS toward glucose-derived OXPHOS [4,5]. Since the heart requires a tremendous amount of energy in the form of ATP to support the demands of the cardiac cycle, it has been suggested that this metabolic rewiring compromises the efficiency of myocardial bioenergetics [5,6]. It is thought that this metabolic rewiring is somewhat inflexible, due to the hypoxic conditions that arise in a hypertrophic muscle [7].

Recent studies [8,9] have explored the therapeutic potential of restoring or enhancing LCFA oxidation to improve cardiac metabolic flux and attenuate hypertrophy. Interventions targeting LCFA transport, mitochondrial uptake, or increasing FAO gene expression with use of peroxisome proliferator-activated receptor (PPAR) agonist drugs have shown promise in both pre-clinical studies and/or clinical trials [10–13]. Yet, the mechanisms of systemic metabolic states and LCFA dynamics in the context of cardiac hypertrophy remain incompletely understood. Further, it is unknown to what extent cardiac metabolism is influenced by whole-body energy balance and lipid profiles, despite the common occurrence of lipotoxicity (accumulation of fatty acid metabolites and intermediates stored in lipid particles) in hypertrophied myocardium [14].

With the growing interest in leveraging metabolism in treating LVH, and a wide range of other cardiovascular diseases (CVDs), a better understanding of the systemic adaptations caused by LVH is necessary. Here, we show that mice with LVH due to the absence of the cardiac-specific regulatory protein, cardiac myosin-binding protein C (cMyBP-C), display a shift in systemic metabolism and depleted energy stores, independent of decreased cardiac output. Our findings suggest that a systemic shift in metabolism is due to the metabolic inflexibility of the heart, requiring other highly metabolic organs, such as liver and skeletal muscle, to compensate for altered cardiac substrate handling.

## Materials and Methods

### Animals

All animal protocols were approved by Case Western Reserve University’s Institutional Animal Care and Use Committee (IACUC), and all experiments were conducted in accordance with the guidelines and regulations set forth in the Animal Welfare Act (AWA) and PHS Policy on Humane Care and Use of Laboratory Animals. All mice were maintained on a 12 hr light/dark cycle in a pathogen-free barrier facility. Male and female *Mybpc3* null (*Mybpc3^−/−^*) mice and SVE129 wild-type controls [15], aged 4-6 months were used in this study.

### Indirect Calorimetry, RER, and Energy Expenditure

Indirect calorimetry (IDC) was used to measure whole-body respiration [16]. This was performed using a Comprehensive Lab Animal Monitoring System (CLAMS; Columbus Instruments). Mice were acclimated to the special caging for 12 hours prior to a 48 hour continuous monitoring period. Mice had *ad libidum* access to 30 g of pelleted food and drinking water for the duration of the experiment. All O_2_ consumption, CO_2_ release, and respirometry exchange ratio (RER) data were normalized to body mass using CLAX software (Columbus Instruments) at the time of data collection. Energy expenditure data were generated using total body weight and average O_2_ consumption during light or dark cycle for each mouse. An ANCOVA test was performed using the JupyterLite Python environment (jupyter.org) with script generated by a large language model (LLM).

### Assessment of Body Composition

Fat stores were measured to estimate mouse body composition. Total body weight was measured for each mouse. Mice were then anesthetized with 5% isopropanol followed by cervical dislocation prior to dissection. The gonadal fat pads — the epidydimal fat pad (EFP) in males or the peri-ovarian adipose tissue (POAT) in females — was harvested and weighed. The gonadal fat pad to total body weight ratio served as an estimate for body composition [17].

### Histology

A histology study was performed to measure organ specific energy stores in cardiac and hepatic tissues. Mice were anesthetized with 5% isoflurane. Hearts and liver were perfused with 1X PBS +1% heparin through the left ventricle followed by perfusion with 4% paraformaldehyde in 1X PBS. Hearts and livers were isolated and further fixed in 4% paraformaldehyde for an additional 24 hours at 4° C. Samples were embedded in paraffin for Hematoxylin & Eosin (H&E), Masson’s trichrome (MT), and Periodic Acid-Schiff (PAS) staining by the Tissue Resources Core at Case Western Reserve University. Snap-frozen heart and liver sections were fixed in 10% formalin and stained with Oil Red O (ORO) by the Tissue Resources Core at Case Western Reserve University. Hematoxylin & Eosin staining was used to study tissue architecture and cell morphology. Masson’s trichrome staining was performed to identify tissue damage due to collagen deposition. Periodic Acid-Schiff staining identified cells with glycogen stores. Slides were scanned at 20 x objective with a Hamastutu Nanozoomer, performed by the Light Microscopy Core at Case Western Reserve University. Heart PAS stains were used to quantify cardiac glycogen stores. Fifteen fields of view were randomly selected from cardiac PAS slides and glycogen stores were quantified within a 600 μM^2^ gated region. Cardiac and hepatic lipid droplets were counted within a 600 μM^2^ gated region in fifteen random fields of view from heart and liver ORO slides, respectively.

### Fasted Glucose and Ketone Measurements

Mice underwent a fasting study to challenge the liver’s ability to regulate in vivo glucose metabolism. Hypoglycemia was induced by placing mice into new cages, singly housed without food for 12 hours (overnight). A tail-snip was performed to express a drop of blood and a hand-held blood glucose and ketone meter was used to measure circulating glucose and ketone levels, as previously described [18].

### Acylcarnitine Analysis

Circulating fatty acid species that tagged for mitochondrial transport by addition of an acyl-CoA group was identified using this clinical mass spectrometry approach. Mice were fasted for 2 hours to normalize postprandial glucose spikes prior to blood serum collection from the inferior vena cava. Serum was sent to the Biomedical Genetics Laboratory at Colorado Children’s Hospital (Aurora, CO). The apparatus consisted of a 1260 Infinity II LC system with a 6470 triple quadrupole mass spectrometer (Agilent Technologies, Palo Alto, CA). Chromatographic separation was achieved on an Acquity UPLC BEH C18 Column (130A pore size, 1.7 μm particle size, 2.1 mm inner diameter × 150 mm length; Waters) held at 50 °C. The method was modified from that developed by Gucciardi et al [19]. The mobile phase was a gradient elution of 0.1 % formic acid in water (A) to 0.1 % formic acid in acetonitrile (B).

The flow rate was 0.4 mL/min, and the gradient was as follows: 0 min 1 %B, 0.1 min 17 %B, 0.28 min 24 %B, 0.35 min 26 %B, 0.80 min 29 %B, 1.71 min 31 %B, 2.96 min 34 %B, 4.50 min 36 %B, 5.44 min 56 %B, 6.37 min 70 %B, 8.01 min 82 %B, 11.30 min 93 %B, 12.50 min 95 %B, 13.50 min 1 %B, 17 min 1 %B. The injection volume was 5 μL and total run time was 17 min per sample. The mass spectrometer was equipped with an ESI source operating in positive mode with gas temp 290 °C, gas flow 5 L/min, nebulizer 35 psi, sheath gas temp 400 °C, sheath gas flow 12 L/min, and capillary voltage 3600 V. Data were acquired in multiple reaction monitoring (MRM) mode. Data were acquired using MassHunter Acquisition 9.0 and processed using MassHunter Quantitative Analysis 10.2.

### High-Resolution Respirometry in Isolated Mitochondria

Mitochondria isolated from left ventricular (LV) and skeletal muscle provided insight into the metabolic flexibility within that tissue. LV tissue (∼45 mg), quadriceps tissue (∼30 mg), or one soleus (∼5 mg) was freshly isolated from wild type or *Mybpc3^−/−^* mice. All tissue was stored and cleaned three times in ice-cold BIOPS buffer containing 10 mM Ca-EGTA (as prepared according to [20]), 0.1 μM CaCl_2_, 20 mM imidazole, 20 mM taurine, 50 mM K-MES, 0.5 mM dithiothreitol, 6.56 mM MgCl_2_, 5.77 mM ATP, and 15 mM phosphocreatine. The tissue was cleaned of ligaments, weighed, and minced using a razor and petri dish on ice. The left ventricle tissue mince was further processed according to [21] to produce freshly isolated mitochondria from the left ventricle (mtLV). The skeletal muscle tissue mince was further processed according to [21] to produce freshly isolated mitochondria from the quadriceps (mtQuad) and soleus (mtSoleus). Intact mitochondria were resuspended in 20 mL/gram of wet tissue mass MiR05 buffer. An Oroboros Oxygraph-2K (O2k) was used to assess Complex I (CI), Complex II (CII), and maximal respiration of isolated mitochondria. 20 μL of freshly isolated mitochondria (approximately 30 μg) was added to 2 mL of O_2_-equilibrated MiR05 buffer in the O2k chambers. Once sample baseline was stable, cytochrome C (0.01 mM) was added to the chamber to assess mitochondrial outer membrane integrity. Malate (1.0 mM), ADP (1.25 mM), pyruvate (5.0 mM), glutamate (5.0 mM), and succinate (10 mM) to stimulate Complexes I & II respiration. Carbonyl cyanide m-chlorophenyl hydrazone (CCCP, 1.0 μM) was added to uncouple the electron transport chain (ETC) and measure maximal oxidation. Finally, rotenone (0.5 μM) and antimycin a (2.5 μM) were added to stop mitochondrial respiration.

### High-Resolution Respirometry of Permeabilized Muscle Fibers

Skeletal muscle – quadriceps and soleus – were isolated from wild type and Mybpc3-/- mice. Each tissue was washed in ice-cold BIOPS buffer, then fibers were teased apart using ultra fine-tip forceps. Tissues were then permeabilized with 5 mg/mL saponin in BIOPS for 20 mins. An Oroboros Oxygraph-2K (O2k) was used to assess Complex I, Complex II, and maximal respiration of permeabilized tissue. 3-5 mg of tissue was added to 2 mL of O_2_-equilibrated MiR05 buffer in the O2k chambers. Once sample baseline was stable, palmitoyl-L-carnitine (0.04 mM) was added to the chamber to supply the tissue with a long chain fatty acid to measure FAO OXPHOS. Malate (0.5 mM), glutamate (10 mM), pyruvate (5 mM), ADP (2.5 mM), cytochrome c (10 μM), and succinate (10 mM) were added to measure Complexes I & II respiration. CCCP (1.0 μM) was added to uncouple the ETC from ATP synthase and measure maximal oxidation. Finally, rotenone (0.5 μM) and antimycin a (2.5 μM) were added to stop mitochondrial respiration.

### Exercise Challenge

To challenge glycolytic capacity and *de novo* glucose synthesis under metabolic stress, mice were run to exhaustion on a single lane isolated rodent treadmill (Columbus Instruments). Exercise naïve mice were acclimated for 10 minutes with belt off and foot shockers on (40 Hz). Data collection began after the acclimation period. Mice walked on the treadmill at 5 m/min, 0% grade for 2.5 mins. The treadmill grade was then increased to 25% for 2.5 minutes at a speed of 5 m/min. The speed was increased from 5 m/min to 14 m/min over the course of 3 mins, then held at 14 m/min until exhaustion. All data were collected at 8 AM. Mice had access to food and water before the exercise challenge.

## Results

### *Mybpc3^−/−^* mice have altered respiration compared to wild type counterparts

Indirect calorimetry was used to determine whether whole-body respiration was different between *Mybpc3**^−/−^*** and wild-type mice. The respiratory exchange ratio (RER), calculated as the ratio of CO_2_ exhaled over O_2_ consumed, is an indicator of whole-body fuel selection, with a ratio of 1.0 indicating pure carbohydrate oxidation and 0.7 indicating pure fat oxidation. Normal mice preferentially oxidize carbohydrates during the dark cycle and shift toward fat oxidation while resting during the light cycle. A moving average of RER in male *Mybpc3**^−/−^*** mice indicated that they had elevated RER values during the 48 hr experiment **(Fig. 1A)**. This was most striking during the light cycle, indicating that *Mybpc3**^−/−^*** mice failed to shift their metabolism toward fat oxidation when less active during the day **(Fig. 1B)**. Similar results were observed in female *Mybpc3**^−/−^*** mice. There was no significant difference in food intake between *Mybpc3**^−/−^*** and wild-type mice of both sexes **(Fig. S1)**.

**Figure 1.**
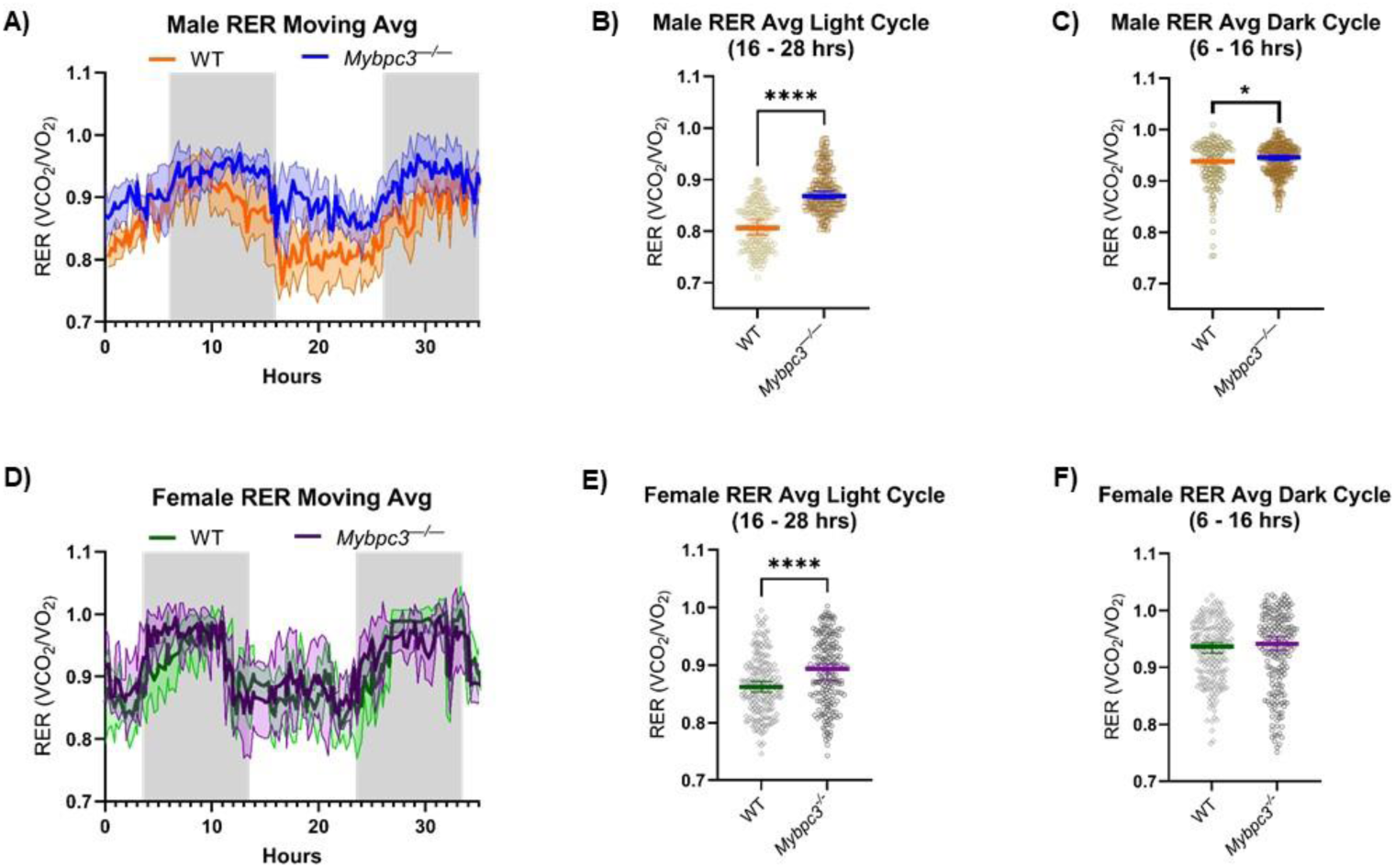
A) IDC data from wild type (N=5) and Mybpc3^-/-^ (N=7) males are represented as a moving average of RER. The average RER of Mybpc3^-/-^ males (AUC = 8.473 ± 0.08646) was higher than wild-type males (AUC = 6.553 ± 0.09763) over the course of 48 hrs. Error bars (shaded region) represent SD. B) Individual RER values from each male mouse cohort was restricted to one light cycle (16 – 28 hrs) for cycle-specific statistical analysis. Mybpc3^-/-^ males (mean = 0.8773) had significantly higher RER readings compared to wild-type males (mean = 0.8079) during the light cycle (unpaired t-test; P<0.0001). Error bars represent SEM. C) Individual RER values from each male mouse cohort was restricted to one dark cycle (6 – 16 hrs) for cycle-specific statistical analysis. Mybpc3^-/-^ males (mean = 0.9398) had significantly higher RER values during the dark cycle compared to wild-type males (mean = 0.9305; unpaired t-test; P=0.0137). Error bars represent SEM. D) Female IDC data from wild-type (N=5) and Mybpc3^-/-^ (N=5) mice are represented as a moving average of RER. The average RER of Mybpc3^-/-^ females (AUC = 7.911 ± 0.1097) was similar to wild-type females (AUC = 7.563 ± 0.09611) over the course of 48 hrs. Error bars (shaded region) represent SD. E) Female Mybpc3^-/-^ mice (mean = 0.8886) had significantly higher RER readings compared to wild-type females (mean = 0.8654) during the light cycle (unpaired t-test; P<0.0001). Error bars represent SEM. F) There was no significant difference in mean RER values between Mybpc3^-/-^ vs wild-type females during the dark cycle (unpaired t-test; P=0.4448). Error bars represent SEM. * = P<0.05; **** = P<0.0001.

To gain further insight into the resting metabolic rate of *Mybpc3**^−/−^*** mice compared to controls, O_2_ consumption was compared between *Mybpc3**^−/−^*** and their wild-type counterparts. A two-way analysis of covariance (ANCOVA) test was performed with genotype and sex as fixed factors, and body mass as a covariate [22]. During the light cycle, there were no significant differences in O_2_ consumption between the two genotypes (F = 0.0862, p = 0.7712) for either sex (F = 0.1400, p = 0711). There were also no significant differences in O_2_ consumption for each genotype (F = 0.0292, p = 0.8655) during the dark cycle in either sex (F = 0.6417, p = 0.4299) (Table 1). This suggested that *Mybpc3**^−/−^*** had a more rigid metabolic flux and were more reliant on glucose oxidation at the whole-body level without influence from differences in O_2_ consumption.

**Table 1.**
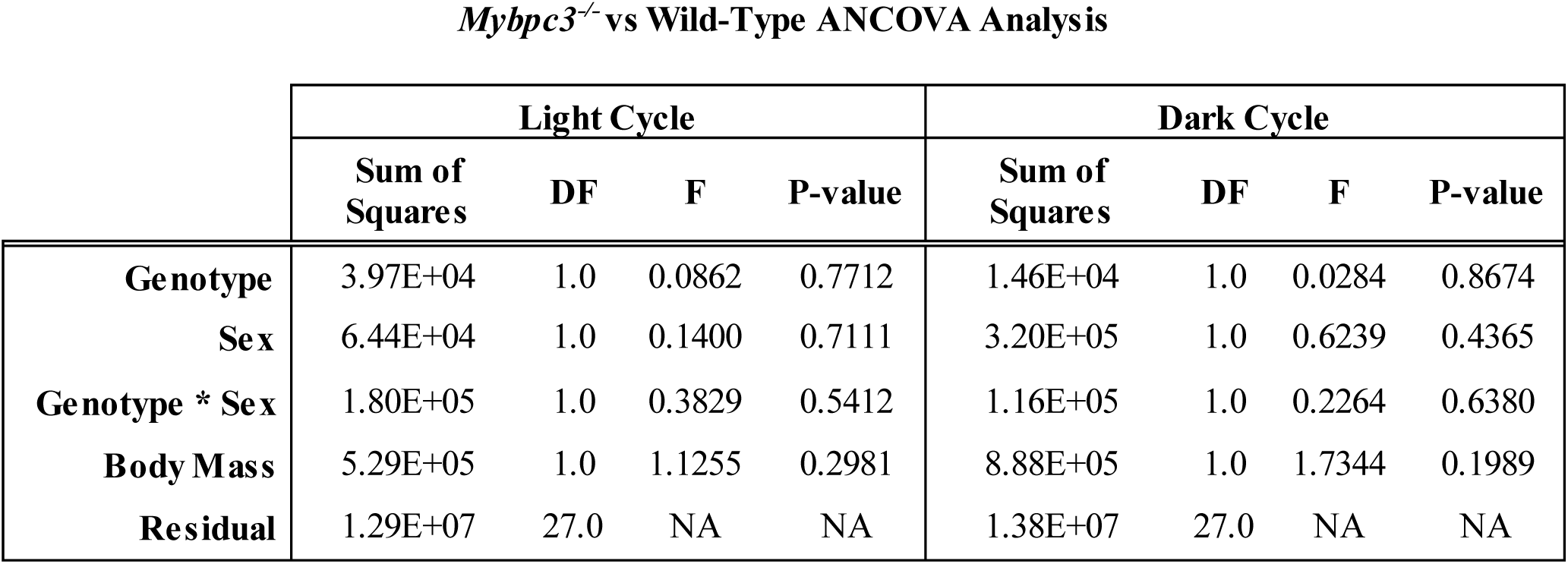
Two-way ANCOVA analysis was performed; assuming that body mass was a covariate of O_2_ Consumption. Genotype, Sex, and Genotype * Sex had no significant effect on O_2_ consumption.

### *Mybpc3^−/−^* mice have altered substrate stores compared to wild-type counterparts

To estimate body composition, the gonadal fat pads of *Mybpc3**^−/−^*** and wild-type male and female mice were isolated, weighed, and normalized to body weight (BW) [17]. Both the average total body weight and average epididymal fat pad (EFP) weight was significantly lower in *Mybpc3**^−/−^*** male mice compared to wild-type counterparts **(Figure 2 A & B)**. When EFP weight was normalized to total BW for each mouse, male *Mybpc3**^−/−^*** mice had a two-fold decrease in EFP / body weight ratio compared to wild-type mice, indicating that *Mybpc3**^−/−^*** mice had significantly less gonadal adiposity compared to wild-type mice **(Figure 2C)**. However, there was no difference in total BW **(Figure 2D)**, periovarian adipose tissue (POAT) **(Figure 2E)**, nor POAT / BW measurements between *Mybpc3**^−/−^*** and wild-type females **(Figure 2F)**.

**Figure 2.**
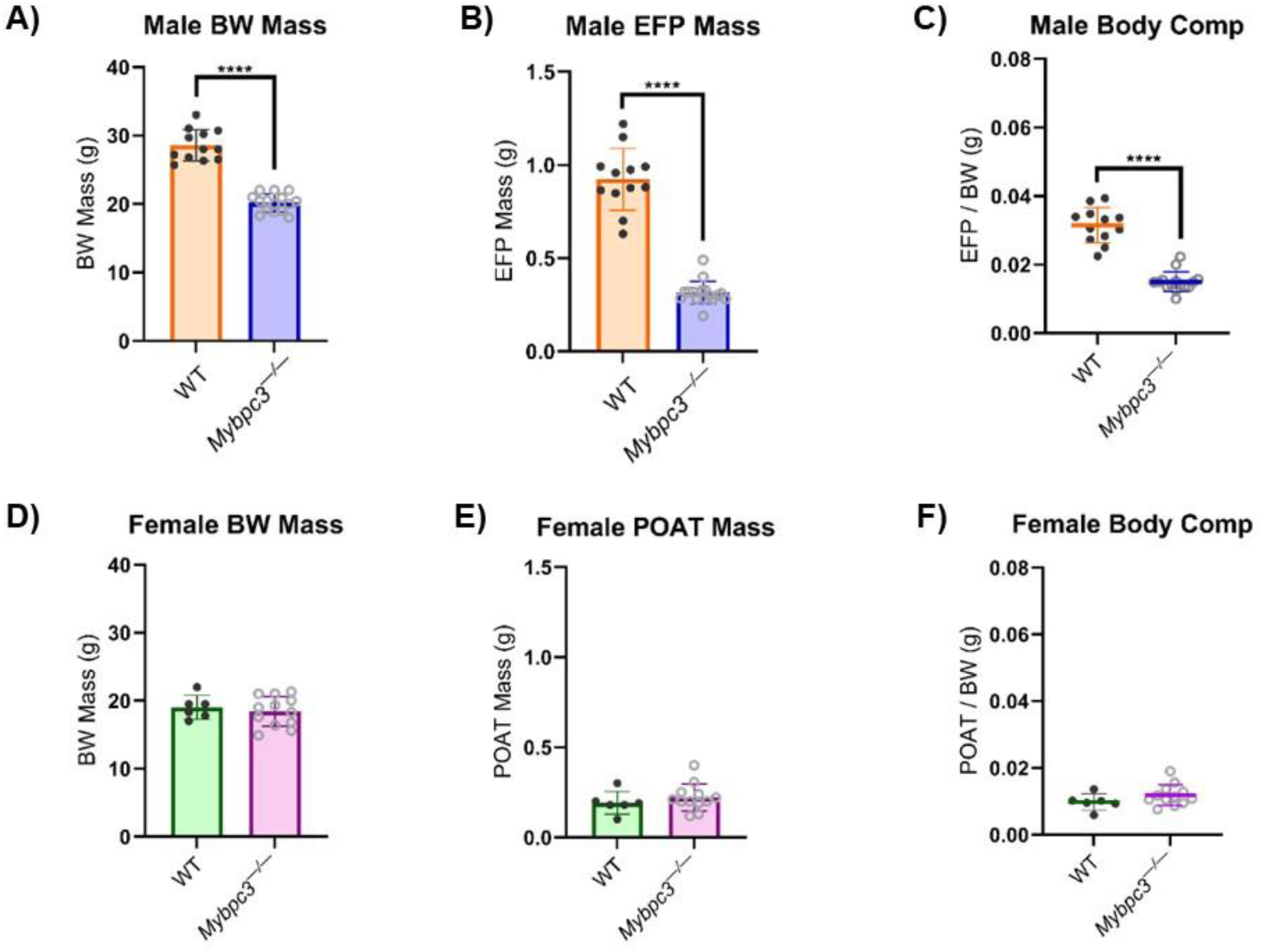
A) Male Mybpc3^-/-^ mice (N=17) had significantly lower BW mass compared to age-matched wild-type males (N=12; unpaired t-test; P<0.0001). B) Male Mybpc3^-/-^ mice (N=17) had significantly lower EFP mass compared to age-matched male wild-type mice (N=12; unpaired t-test; P<0.0001). C) Male EFP weights were normalized to BW to calculate an EFP / BW ratio to serve as an estimation of body composition. An unpaired t -test showed male Mybpc3^-/-^ mice had a significantly lower EFP / BW ratio compared to wild type males (unpaired t-test; P<0.0001). D) There was no difference in female Mybpc3^-/-^ mice (N=12) BW mass compared to age-matched wild-type females (N=6; unpaired t-test; P=0.5803). E) There was no significant difference between female Mybpc3^-/-^ mice (N=12) POAT mass compared to age-matched female wild-type mice (N=6) POAT mass (unpaired t-test; P=0.4091). F) Female POAT weights were normalized to BW to calculate a POAT / BW ratio to serve as an estimation of body composition. An unpaired t-test showed female Mybpc3^-/-^ and female wild-type mice had no significant difference in POAT / BW ratio (unpaired t-test; P=0.4091). **** = P<0.0001. All error bars represent SD.

Given the increased RER and lack of adipose tissue in male *Mybpc3**^−/−^*** mice, we wanted to investigate whether *Mybpc3**^−/−^*** mice had access to the fatty acids, stored as triglycerides in lipid droplets, that drive FAO OXPHOS in oxidative tissues. A histology study with Oil Red O (ORO) staining was performed to measure lipid droplet accumulation in cardiac **(Figure 3)** and liver **(Figure 4)** tissue collected from *Mybpc3**^−/−^*** and wild-type mice. The wild-type mice had significantly less lipid droplet accumulation in cardiac tissue compared to *Mybpc3**^−/−^*** tissue **(Figure 3E)**, suggesting this model of LVH had lipotoxicity, consistent with clinical data and other animal models [23–25]. *Mybpc3**^−/−^*** mice also had increased lipid droplet accumulation in hepatic tissue **(Figure 4E)**, suggesting a fatty liver phenotype [26]. Quantitative analysis of Periodic acid-Schiff (PAS) staining showed that *Mybpc3**^−/−^*** mice had significantly more glycogen positive cardiomyocytes compared to wild-type mice **(Figure 3F)**. Whole-slide scans of liver PAS stains suggested fewer glycogen-rich regions in *Mybpc3**^−/−^*** livers compared to wild-type livers **(Figure S2)**.

**Figure 3.**
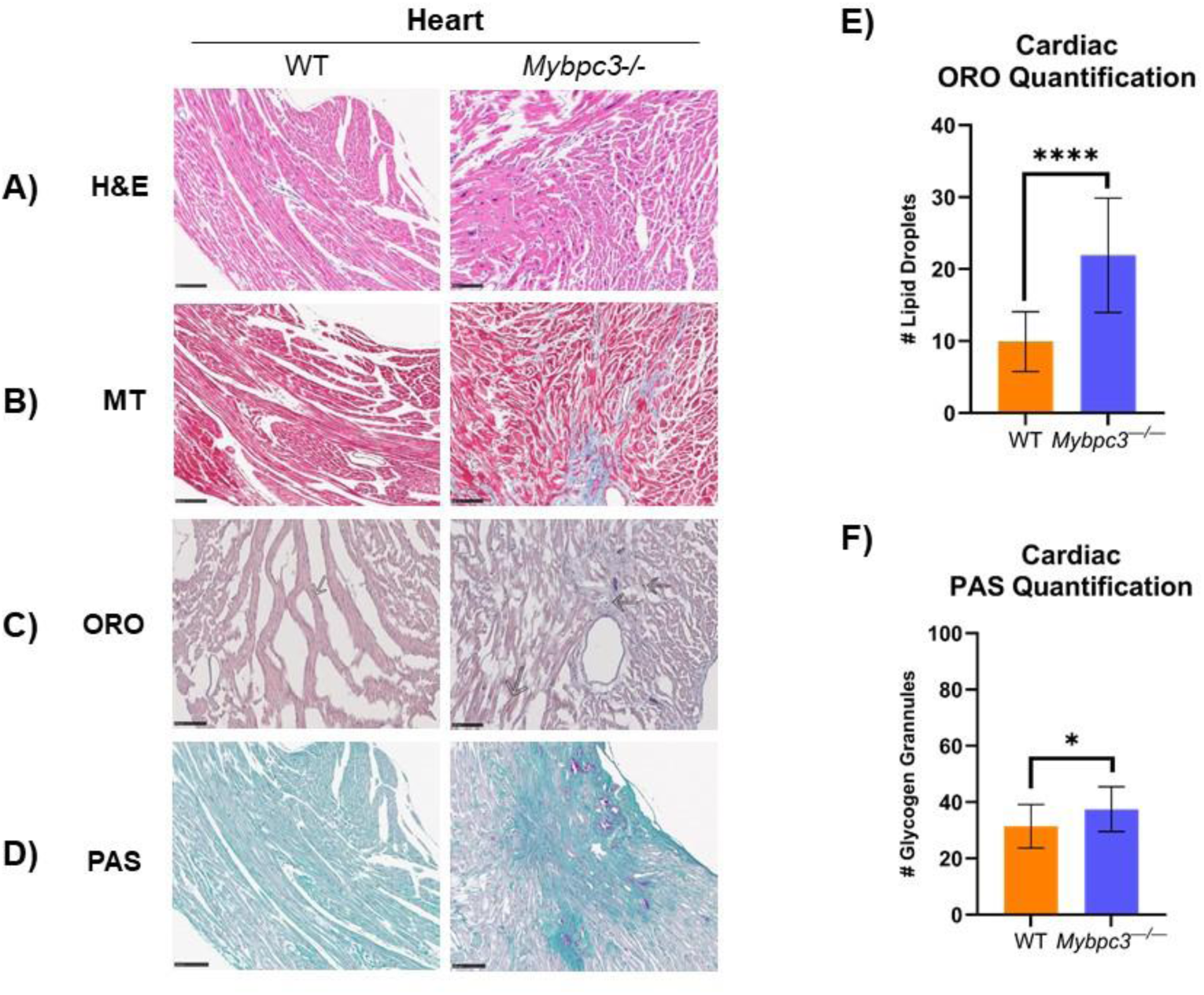
Histological analysis was performed in cardiac tissue collected from male Mybpc3^-/-^ and wild-type mice. A) H&E staining shows Mybpc3^-/-^ mice have differences in cardiomyocyte size and organization. B) MT staining shows that Mybpc3^-/-^ mice have clear, large regions of collagen deposition. C) ORO staining shows that Mybpc3^-/-^ cardiac tissue has accumulation of large lipid droplets (as indicated by arrows). D) PAS staining indicated glycogen-positive cardiomyocytes in both wild-type and Mybpc3^-/-^ cardiac tissues. Scale bar represents 100 μM; N=3 for each group. E) The number of lipid droplets per 600 μM in Mybpc3^-/-^ mouse (N=15) cardiac tissue was significantly higher compared to wild-type mice (N=15; unpaired t-test; P<0.0001). F) The number of glycogen granules per 600 μM was significantly higher in Mybpc3^-/-^ (N=15) cardiac tissue compared to wild-type mice (N=15; unpaired t-test; P=0.0460). * = P<0.05; **** = P<0.0001. All error bars represent SD.

**Figure 4.**
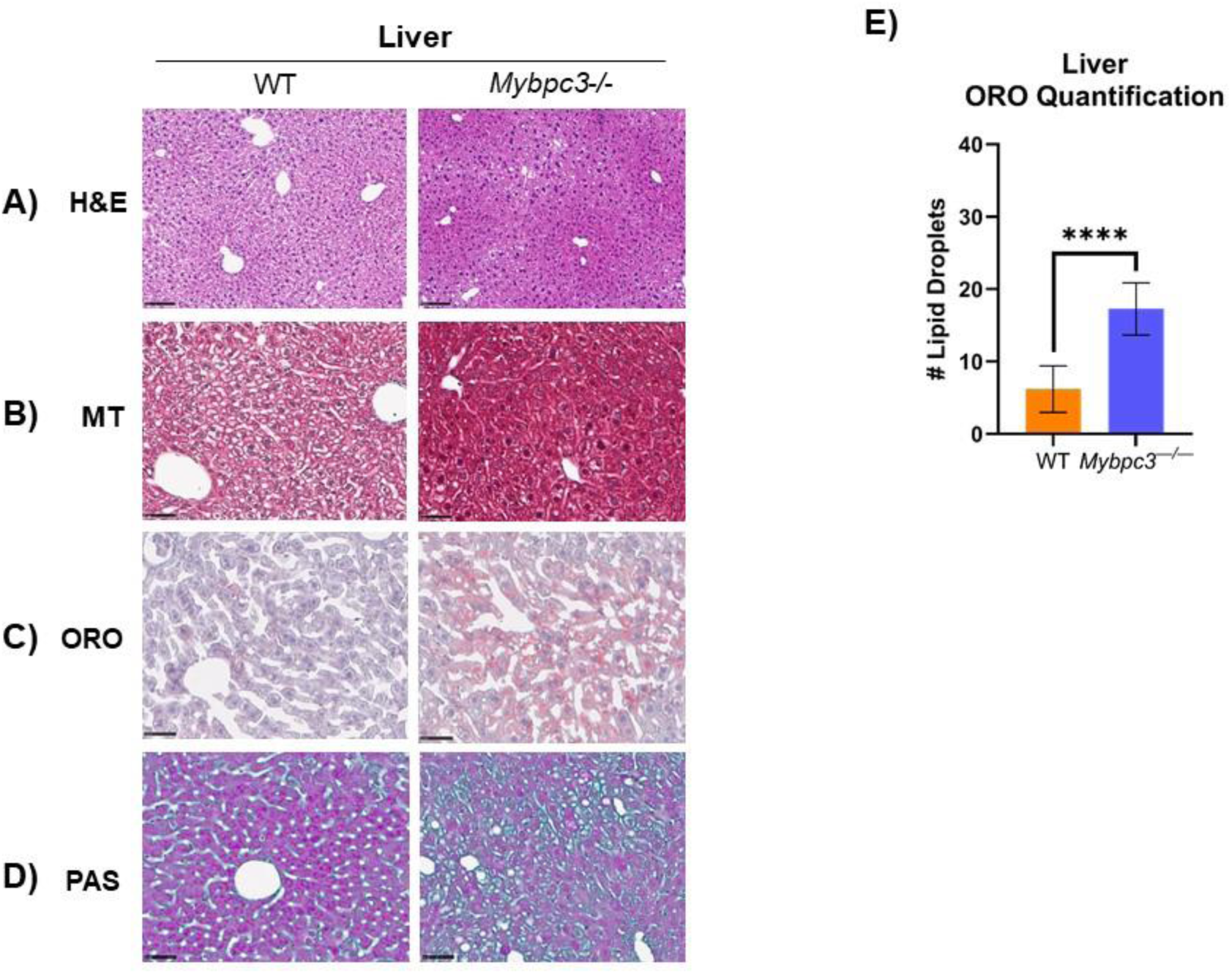
A) Histological analysis was performed in liver tissue collected from male Mybpc3^-/-^ and wild-type mice. H&E staining shows no clear differences in hepatocyte size nor organization. MT staining shows that Mybpc3^-/-^ mice have clear, large regions of collagen deposition. ORO staining shows that Mybpc3^-/-^ hepatic tissue has accumulation of large lipid droplets. PAS staining indicated glycogen-positive cardiomyocytes in both wild-type and Mybpc3^-/-^ cardiac tissues. Scale bar represents 100 μM; N=3 for each group. B) Mybpc3^-/-^ mice (N=15) had significantly more lipid droplets in liver per 600 μM compared to wild-type mice (N=15; P<0.0001). **** = P<0.0001. Error bars represent SD.

Hematoxylin & Eosin (H&E) and Masson’s Trichrome (MT) staining were also performed in these tissues to assess tissue architecture, hypertrophy, and tissue damage **(Figures 3 & 4)**. H&E stains of cardiac tissue showed dysmorphic, unorganized cardiomyocytes in the *Mybpc3**^−/−^*** mice, due to the LVH phenotype **(Figure 3A)**. MT staining showed collagen deposition and development of fibrotic tissue in *Mybpc3**^−/−^*** males, as expected **(Figure 3B)**. H&E and MT staining did not uncover any tissue damage in *Mybpc3**^−/−^*** liver **(Figure 4A)**, despite lipid droplet accumulation **(Figure 4C & 4E)** and potential reduction of glycogen stores **(Figure D)**.

### Circulating metabolites differ between *Mybpc3^−/−^* and wild-type males

Since both light and dark cycle RER measurements, and body composition estimates were significantly different in males, but not females, we focused further metabolic flux experiments on male mice. To determine whether *Mybpc3**^−/−^*** mice had altered circulating metabolites potentially causing altered substrate stores in heart and liver, blood glucose and ketone levels were measured before and after a 12 hour fasting challenge. Blood glucose levels in wild-type animals significantly decreased post-fast, as expected. *Mybpc3**^−/−^*** male glucose levels were significantly lower than wild type mice before and after the 12 hr challenge, suggesting inflexibility of carbohydrate metabolism **(Figure 5A)**. *Mybpc3**^−/−^*** males had significantly higher blood ketone levels before and after the fasting challenge **(Figure 5B)**. Mean blood ketone levels almost tripled from 0.35 mmol/L in a fed state to 0.90 mmol/L in a fasted state in wild-type mice. However, *Mybpc3**^−/−^*** mice had mean blood ketone levels around 2.0 mmol/L in both a fed and fasted state; reaching blood ketone levels typically observed in mouse models of ketoacidosis [27,28].

**Figure 5.**
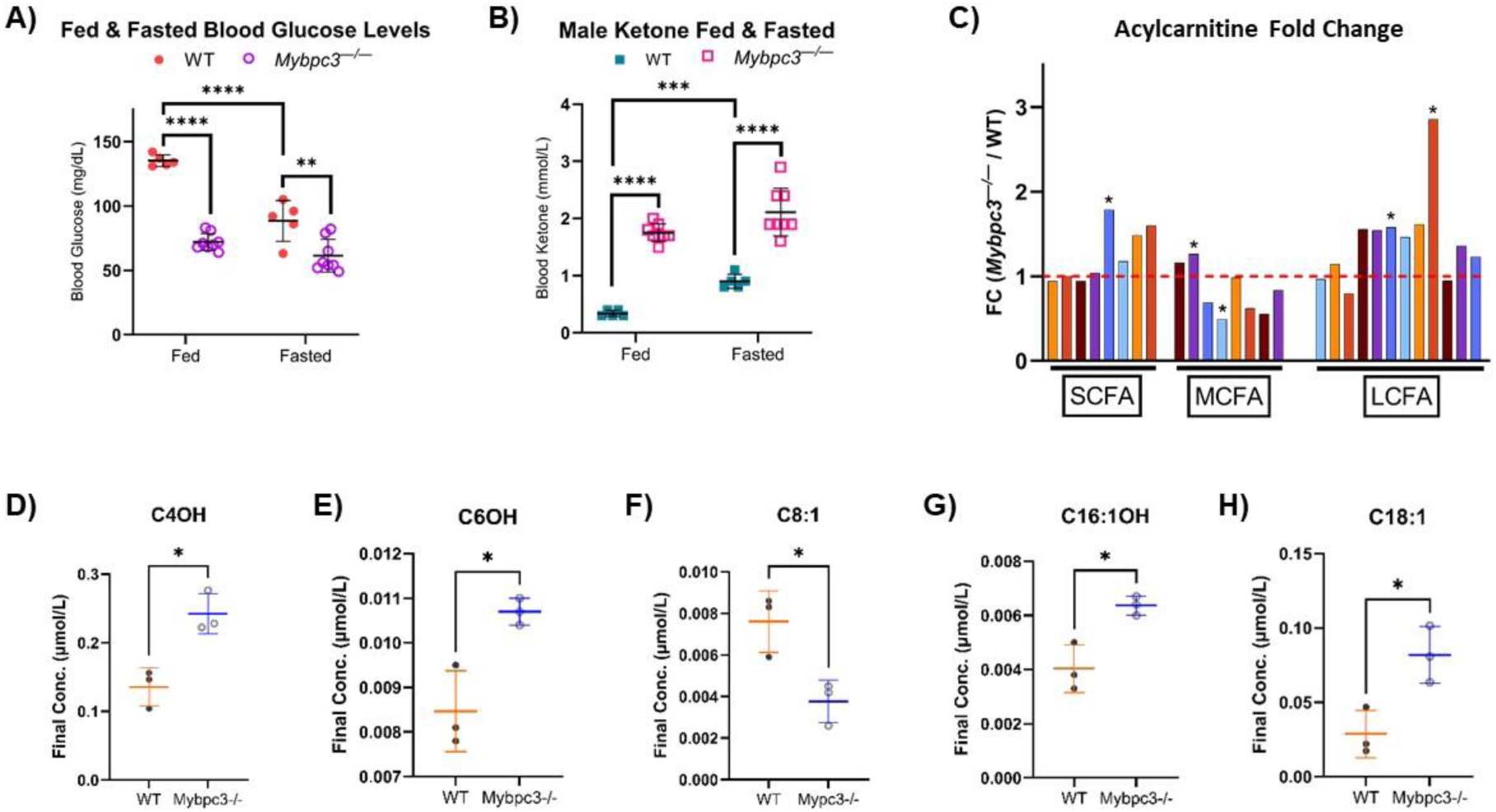
A) Two-way ANOVA mixed-effects analysis was performed to determine the effect of fasting on blood glucose levels in Mybpc3^-/-^ (N=8) vs wild-type (N=8) male mice. Overall, fasting only decreased blood glucose levels in wild-type mice. Fed wild-type mice had significantly higher blood glucose levels compared to Mybpc3^-/-^ mice (2-way ANOVA; P<0.0001). B) Male Mybpc3^-/-^ had significantly higher blood ketone levels compared to wild-type mice in both a fed (2-way ANOVA; P<0.00001) and fasted (2-way ANOVA; P<0.00001) state. Wild-type mice had a significant increase in blood ketone levels in a fasted state compared to a fed state (2-way ANOVA; P<0.0001). C) Fold change (FC) was calculated for each acyl-CoA species by dividing mean Mybpc3^-/-^ concentration by mean wild - type concentration. Values over 1.0 indicate an increase in Mybpc3^-/-^ mice; values below 1.0 indicate a decrease in Mybpc3^-/-^ mice. P-values were calculated using the mean of each genotype to determine significant increases/decreases. D – H) All acylcarnitine chain lengths with significant differences in mean concentrations are shown. * = P<0.05; ** = P<0.005; *** = P<0.0005; **** = P<0.0001. Error bars represent SD.

Circulating glucose and ketone data indicated that *Mybpc3**^−/−^*** mice were in metabolic duress at the whole-body level. To better understand how this affected metabolism of fatty acids, an acylcarnitine profiling study was performed in blood serum. The acylcarnitine profile is a well-known biomarker of FAO flux *in vivo*. We found that the biggest difference in acylcarnitines was a significant, three-fold increase in C18:1-carnitine in *Mybpc3**^−/−^*** mice **(Fig 5H)**. In fact, there was a slight elevation in most long chain-carnitine species **(Figure 5C)**, suggesting that *Mybpc3**^−/−^*** mice were not only more reliant on glucose catabolism and oxidation for energy production, but that FAO-linked OXPHOS was inefficiently using available LCFAs after first-pass FAO.

The elevated ketones shown earlier in *Mybpc3**^−/−^*** mice indicated higher liver long chain-FAO, as FAO is the source of mitochondrial acetyl-CoA during ketogenesis. Yet, the serum acylcarnitine profile indicated inefficient FAO at the whole-body level. It has been shown that the serum acylcarnitine profile primarily reflects cardiac muscle FAO, and secondarily skeletal muscle FAO [29]. Muscle is also the major contributing factor to RER measurements at rest and during exercise [30], which is further influenced by muscle fiber type [31]. To determine whether the apparent metabolic inflexibility in *Mybpc3**^−/−^*** mice reflects changes in muscle physiology, exercise naïve mice were given an acute exercise challenge. Indirect calorimetry was used to monitor RER during the initial 15 mins of treadmill running, with glucose and ketone body levels measured pre- and post-exhaustion. *Mybpc3**^−/−^*** mice had significantly higher RER and blood ketone levels pre-exhaustion, as described earlier. Wild-type mice ran significantly further than *Mybpc3**^−/−^*** mice, indicating that LVH caused an exercise intolerance phenotype . *Mybpc3**^−/−^*** mice did have similar RER to wild-type mice as the exercise challenge progressed past the “aerobic” portion **(Figure 6A**). However, *Mybpc3**^−/−^*** mice post-exhaustion had significantly higher glucose levels compared to wild-type mice **(Figure 6B)**. Ketone levels were equal pre- and post-exhaustion in both *Mybpc3**^−/−^*** and wild-type mice. Although, *Mybpc3**^−/−^*** mice did have significantly higher blood ketone levels pre- and post-exhaustion **(Figure 6C)**. Hypoxia-inducible factor 1-alpha (Hif-1α) is a known regulator of glycolysis and has been shown to be elevated in *Mybpc3**^−/−^*** heart due to the LVH phenotype [32]. To test whether a similar mechanism extends to skeletal muscle, immunoblotting was performed to detect Hif-1α. Densitometric analysis indicated no differences in Hif-1α protein levels between the two genotypes **(Figure 6E & 6F)**.

**Figure 6.**
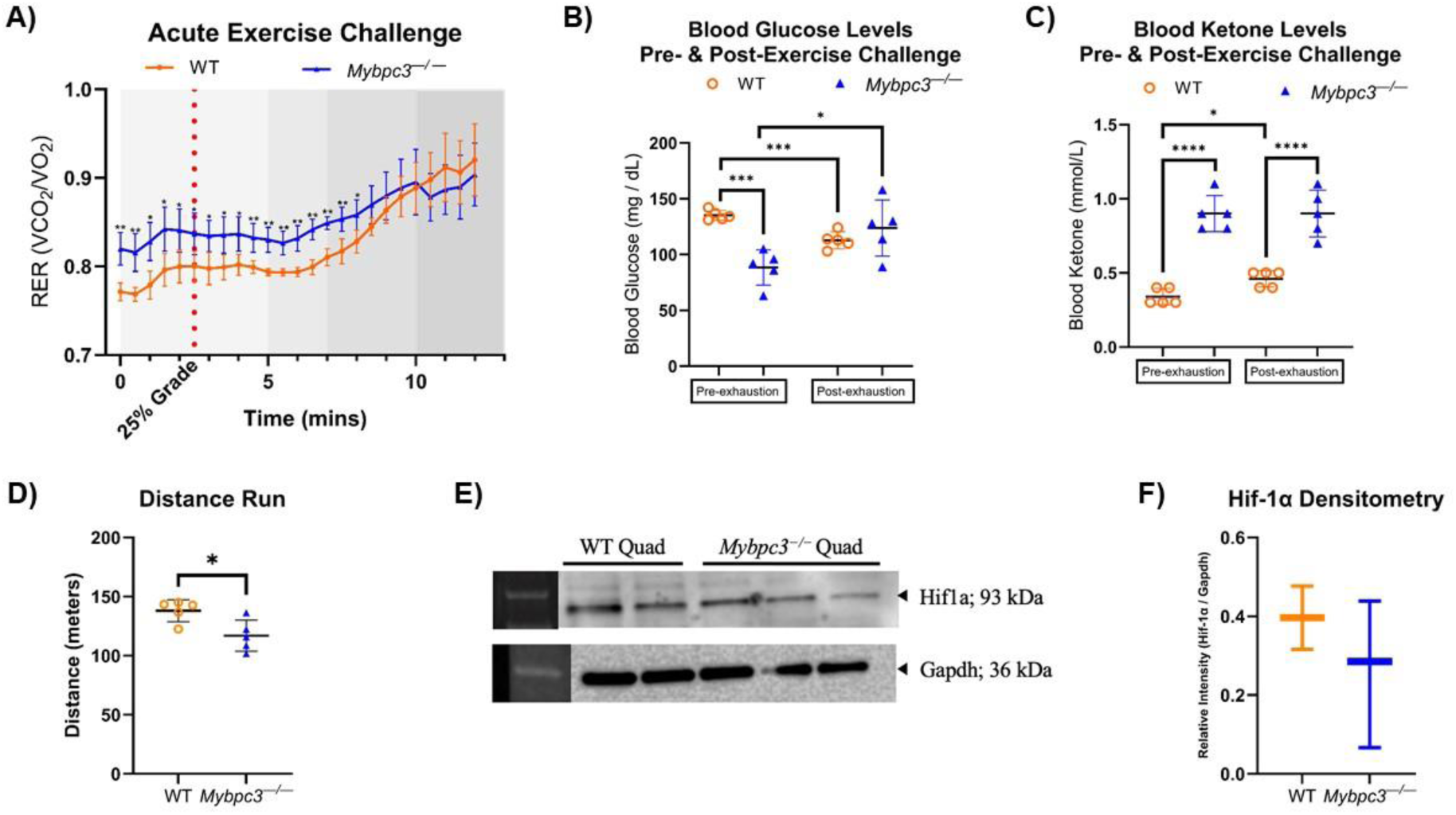
A) RER values within each genotype were averaged and plotted as a moving RER average against time. Unpaired t-tests to compare wild-type and Mybpc3^-/-^ mean at each time point (X-axis) were performed. B) Blood glucose levels measured pre- and post-exhaustion were plotted for Mybpc3^-/-^ male mice (N=5) and wild-type mice (N=5). A paired t-test showed that wild-type mice had a significant decrease in blood glucose levels post-exhaustion (P=0.0009). A paired t-test showed that Mybpc3^-/-^ mice (N=5) had a significant increase in blood glucose levels post-exhaustion (P=0.0429). An unpaired t-test showed that Mybpc3^-/-^ mice had significantly lower blood glucose levels than wild-type mice pre-exhaustion (P=0.0002). C) Blood ketone levels measured pre- and post-exhaustion were plotted for Mybpc3^-/-^ male mice (N=5) and wild-type mice (N=5). Mybpc3^-/-^ mice had significantly higher blood ketone levels pre-exhaustion compared to wild-type mice (P<0.0001). Mybpc3^-/-^ mice had significantly lower blood ketone levels compared to wild-type mice post-exhaustion (P<0.0001). Wild-type mice had significantly higher blood ketone levels post-exercise challenge (P=0.0327). D) Mybpc3^-/-^ mice ran a significantly shorter distance compared to wild-type mice (unpaired t-test; P = 0.0192). E) Immunoblot data showed that Hif-1α was present at low levels in both Mybpc3^-/-^ and wild-type quadriceps. Anti-Gapdh was used as a control. F) Densitometric analysis of Hif-1α band intensity normalized to Gapdh band intensity showed no significant difference in protein levels (P=0.3976) between Mybpc3^-/-^ mice (N=3) vs wild-type mice (N=2). * = P<0.05; ** = P<0.005; *** = P<0.0001; **** = P<0.00001. All error bars represent SD.

### *Mybpc3^−/−^* males have decreased OXPHOS capacity in heart and skeletal muscle compared to wild-type males

Mitochondria isolated from the LV (mtLV) of *Mybpc3**^−/−^*** mice have significantly decreased oxygen consumption compared to wild-type mtLV with the supplementation of oxidative substrates pyruvate, glutamate, and succinate **(Figure 7A)**. Both NADH-dependent complex CI, FADH_2_-activated complex CII, and maximal OXPHOS capacity were significantly reduced in *Mybpc3**^−/−^*** mtLV **(Figure 7B)**. However, State III respiration was not significantly different. This suggested that the proton gradient, which is in essence potential energy within the inner mitochondrial membrane (IMM) in the form of H^+^, could be disrupted in *Mybpc3**^−/−^*** mouse mtLV.

**Figure 7.**
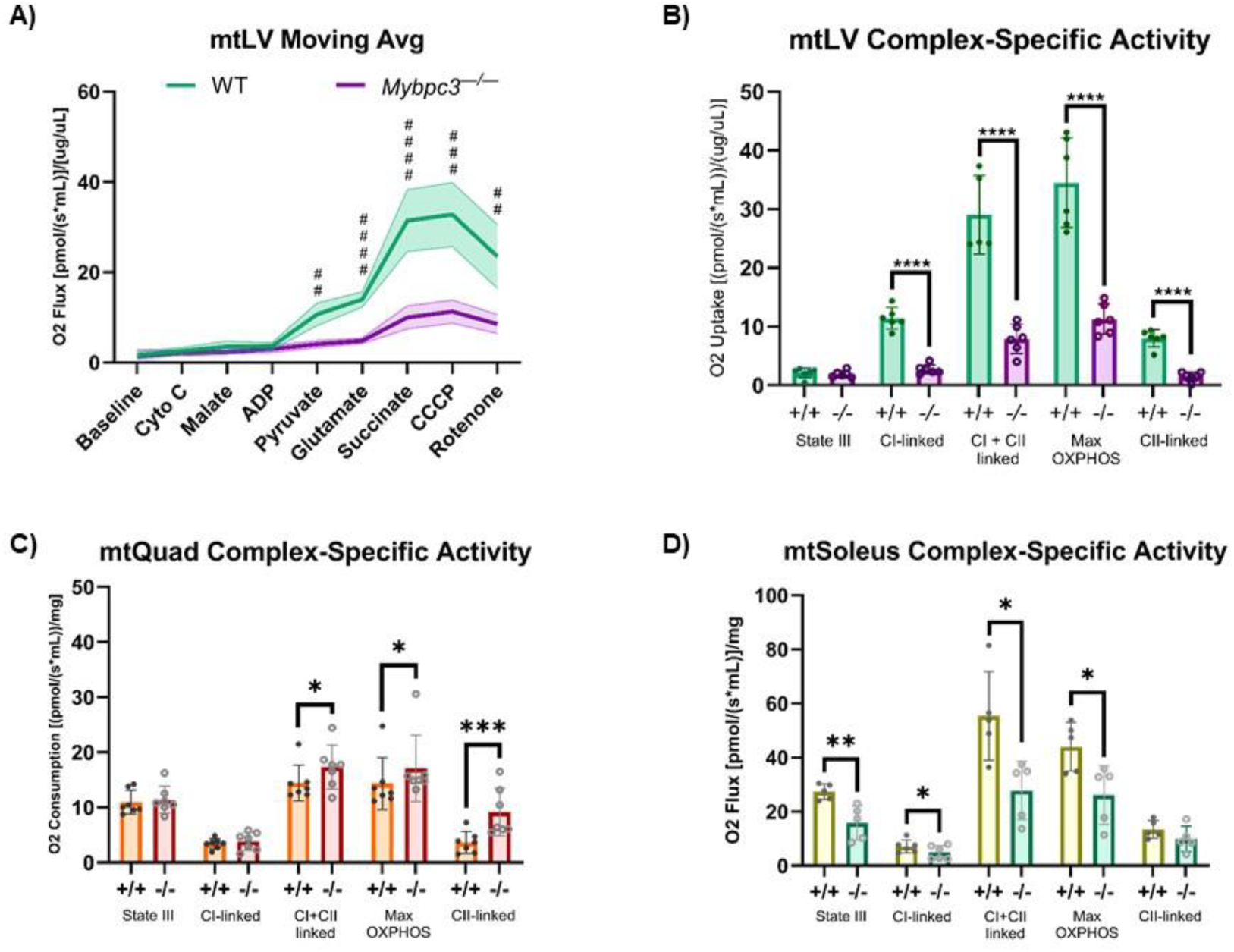
A) The mean respiration of wild-type mtLV (N = 6) and Mybpc3^-/-^ mtLV (N = 6) is represented as a moving average in response to acute treatment of various metabolic substrates. AUC analysis showed that wild type mt LV had an AUC value of 111.20±8.23 compared to mtLV Mybpc3^-/-^ AUC value 42.48±3.08. Unpaired t-test analysis showed that Mybpc3^-/-^ mtLVs had significantly lower mitochondrial respiration in response to pyruvate (Q=0.000111), glutamate (Q<0.000001), succinate (Q=0.000090), CCCP (P=0.000090), and rotenone (Q0.000632) treatment. B) Complex-specific activity was calculated in wild type (N = 6) and Mybpc3^-/-^ (N = 6) mtLV. Mybpc3^-/-^ mtLV had significantly reduced CI-linked (P<0.0001), CI + CII-linked (P<0.0001), Max OXPHOS (P<0.0001), and CII-linked (P<0.0001) compared to wild type mtLV. C) Complex-specific activity was calculated from mean oxygen consumption in mtQuad isolated from wild type mice (N = 7) and Mybpc3^-/-^ mice (N = 7) quadriceps. Unpaired t-tests showed that mtQuad from Mybpc3^-/-^ mice was significantly increased in CI+CII-linked (P=0.0111), Max OXPHOS (P=0.0469), and CII-linked (P=0.0030) compared to wild-type mtQuad. D) Complex-specific activity was calculated from mean oxygen consumption in mtSoleus isolated from wild-type mice (N = 5) and Mybpc3^-/-^ mice (N = 5) soleus. Unpaired t-tests showed that mtSoleus from Mybpc3^-/-^ mice had significantly decreased State III (P=0.0060), CI-linked (.0430), CI+CII-linked (P=0.0140), and Max OXPHOS (P=0.0222) compared to wild-type mtSoleus.

Given that we found systemic metabolism was significantly different between *Mybpc3**^−/−^*** and wild-type males, it is possible that skeletal muscle mitochondria could have differences in metabolism indirectly caused by LVH. We found that mitochondria isolated from the quadriceps (mtQuad), which is a glycolytic, fast-twitch muscle, had no differences in State III nor CI-linked activity in *Mybpc3**^−/−^*** activity compared to wild-type mtQuad. However, CII-linked, CI-+ CII-linked (likely due to increased CII activity), and maximal OXPHOS were all significantly higher in *Mybpc3**^−/−^*** mtQuad **(Figure 7C)**. Similar to mtLV, mitochondria isolated from slow-twitch, FAO-dependent soleus tissue (mtSoleus) in *Mybpc3**^−/−^*** mice had significantly lower CI-linked, CI- + CII-linked, and maximal OXPHOS capacity compared to wild-type mtSoleus **(Figure 7D)**.

### Long chain-fatty acid supplementation ameliorates decreased OXPHOS capacity in *Mybpc3^−/^*^−^ male skeletal muscle in vitro

Our finding that reduced OXPHOS capacity in a highly metabolic tissue outside the heart was surprising. Moreover, that this reduced OXPHOS capacity seemed specific to tissue that typically rely on FAO-mediated OXPHOS, as soleus tissue does [33]. We hypothesized that exogenous supplementation of a long-chain fatty acid (LCFA), palmitoyl-L-carnitine would reverse the reduced mitochondrial capacity in mtLV and mtSoleus. To test this, muscle fibers from quadriceps and solei were collected and permeabilized with saponin as previously described [34]. This facilitated LCFA uptake independent of fatty acid transporters. Treatment of permeabilized muscle fibers from quadriceps (pQuad) did not appear to affect FAO-mediated OXPHOS between *Mybpc3**^−/−^*** and wild-type pQuad. However, significantly increased CCCP response and max OXPHOS data indicated that maximal OXPHOS capacity was slightly increased in *Mybpc3**^−/−^*** pQuad after treatment of the tissue fibers with palmitoyl-L-carnitine **(Figure 8A & B).** Supplementation of palmitoyl-L-carnitine to pSoleus increased oxygen flux in *Mybpc3**^−/−^*** solei by almost two-fold **(Figure 8C)**. Furthermore, linked complex activities were all significantly increased in *Mybpc3**^−/−^*** pSolei, with a three-fold increase in both CI + CII-linked OXPHOS and CII-linked OXPHOS **(Figure 8D)**. Maximum OXPHOS capacity was also significantly higher in pSoleus from *Mybpc3**^−/−^***mice.

**Figure 8.**
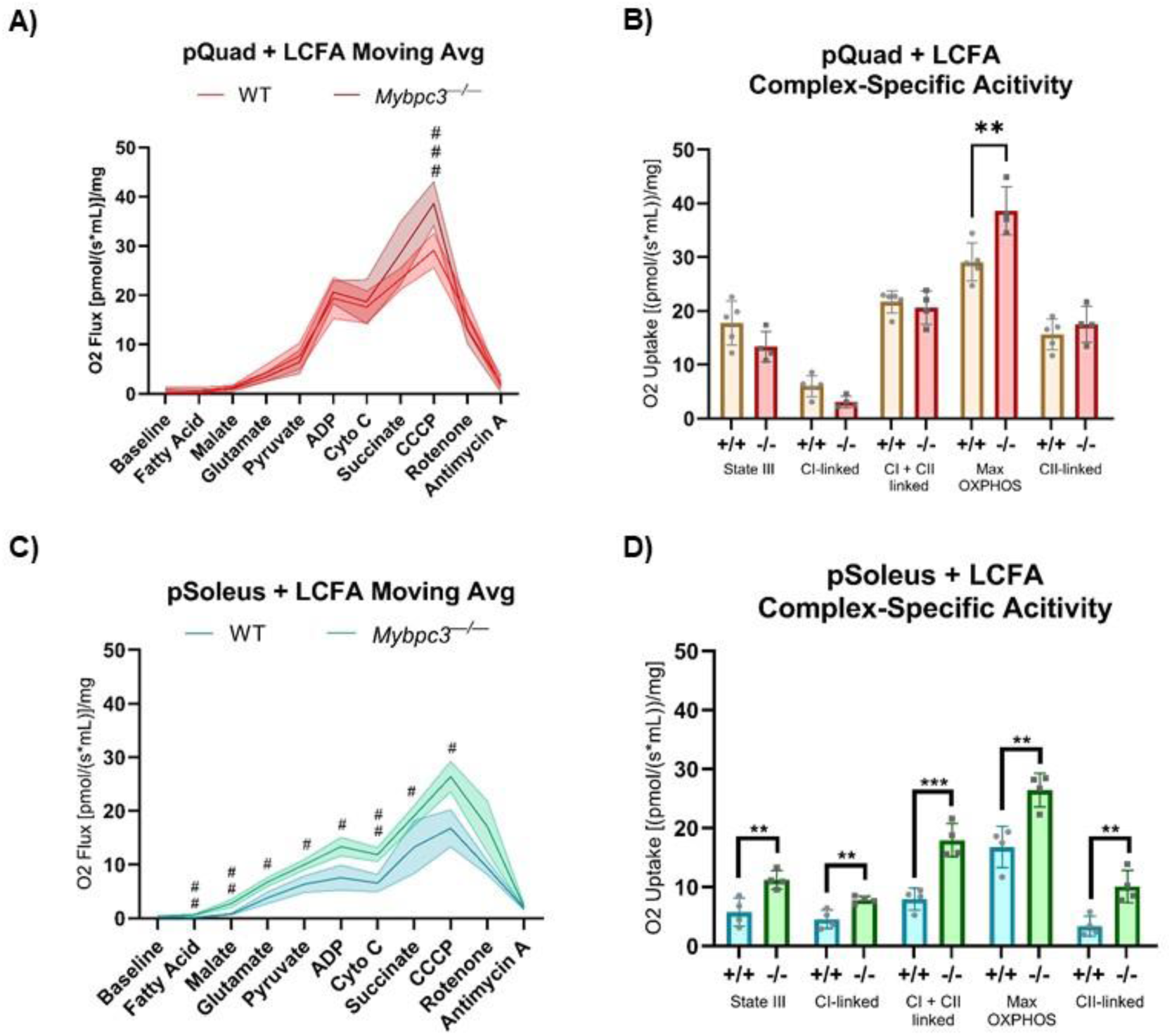
A) Metabolic flux is represented as oxygen flux in response to acute treatment of various substrates in permeabilized quadriceps muscle fibers (pQuad) collected from Mybpc3^-/-^ mice (N = 4) and wild-type mice (N = 5). AUC analysis indicated that wild-type and Mybpc3^-/-^ pQuad had similar area values, with 119.5±5.664 and 132.5±7.173, respectively. Mybpc3^-/-^ pQuad had a significantly higher response to CCCP (Q < 0.0001) compared to wild-type mtQuad. Error bars represent SD. B) State III, maximum OXPHOS capacity, and complex-specific activity were calculated for pQuad collected from Mybpc3^-/-^ and wild-type mice with treatment of a long chain-fatty acid (LCFA). Mybpc3^-/-^ pQuad (N=4) had significantly higher oxygen uptake only at maximal OXPHOS compared to wild-type pQuad (N=5; P = 0.0089). C) Metabolic flux is represented as oxygen flux in response to acute treatment of various substrates, including a LCFA, in permeabilized soleus muscle fibers (pSoleus). AUC analysis showed that the total peak area in Mybpc3^-/-^ pSoleus was higher than wild-type pSoleus, with 109.1±4.557 and 65.7±5.007, respectively. Mybpc3^-/-^ pSoleus samples had significantly higher oxygen influx after fatty acid (Q=0.0061), malate (Q = 0.0061), glutamate (Q = 0.0085), pyruvate (Q = 0.0091), ADP (Q = 0.0091), cytochrome c (Q = 0.0061), and CCCP (Q = 0.0085) treatment. Error bars represent SD. D) State III, maximum OXPHOS capacity, and complex-specific activity were calculated for pSoleus collected from Mybpc3^-/-^ and wild-type mice with treatment of a long chain-fatty acid (LCFA). State III (P = 0.0087), CI-linked (P=0.0083), CI- + CII-linked (P = 0.0010), maximum OXPHOS (P = 0.0052), and CII-linked (P = 0.0059) were all significantly increased in Mybpc3^-/-^ mice (N = 4) compared to wild-type mice (N = 4). Error bars represent DS. # = Q<0.01; ## = Q<0.001; ### = Q<0.0001. * = P<0.05; ** = P<0.005; *** = P<0.0005.

### Exogenous supplementation of LCFAs bypasses lipid storage in *Mybpc3^−/−^* mice

Since we observed a significant increase in mitochondrial respiration in *Mybpc3**^−/−^*** soleus tissue, we hypothesized that systemic *Mybpc3**^−/−^*** metabolism would increase as a result of exogenous LCFA supplementation in the form of a high fat diet (HFD; >60% kcal from lard). *Mybpc3**^−/−^*** and wild-type controls of both sexes were fed a HFD for 10 weeks to test whether exogenous LCFA supplementation could affect the differences in body composition, metabolic deficits, and substrate handling *in vivo*. Both male and female Mybpc3-/- mice had significantly lower mean BWs **(Figure 9A)** and GFP masses **(Figure 9B)** compared to wild-type mice. Although both male and female *Mybpc3**^−/−^*** mice on a HFD had significantly reduced GFP adiposity compared to wild-type mice **(Figure 7C)**, intracellular lipid stores appeared to change as a result of HFD. ORO staining of wild-type cardiac tissue indicated accumulation of large lipid droplets, which were not present in Mybpc3-/- tissue samples **(Figure 9D)**. Clear, large lipid droplets were visible in H&E, MT, ORO, and PAS stains of wild-type liver tissues. However, *Mybpc3**^−/−^*** mice didn’t appear to have similar clear, large lipid droplets visible in H&E, MT, ORO, or PAS stains **(Figure 9E)**. Both *Mybpc3**^−/−^*** and wild-type mice had glycogen-rich and glycogen-depleted regions in hepatic tissue **(Figure S5)**. Despite these differences in lipid accumulation, *Mybpc3**^−/−^*** on a HFD presented with similar pathogenic changes in tissue architecture and collagen deposition as *Mybpc3**^−/−^*** mice on a standard lab diet, indicating that the HFD did not rescue LVH pathogenesis **(Figure 9D)**.

**Figure 9.**
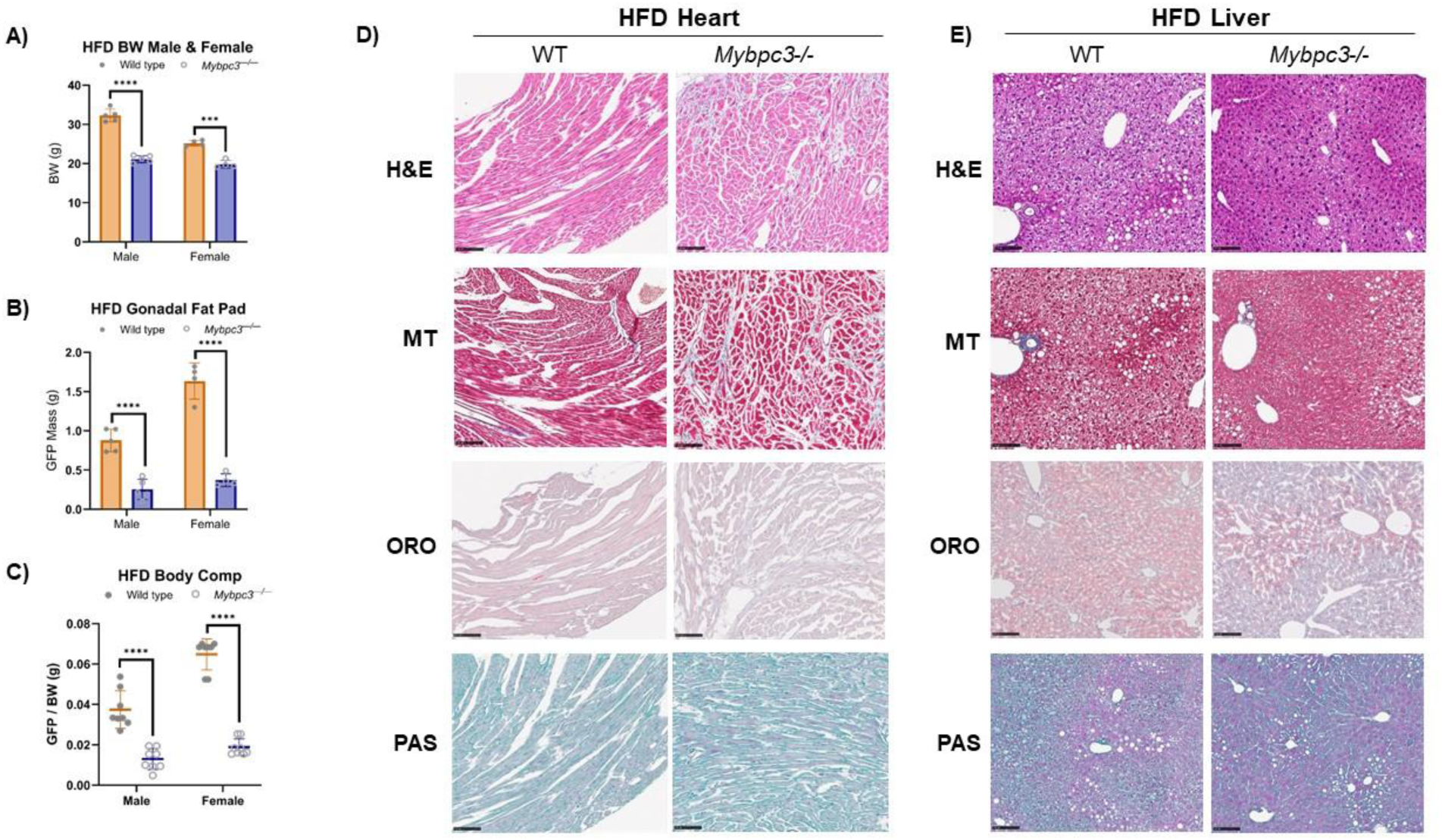
Mybpc3^-/-^ mice and wild-type mice of both sexes were fed a HFD for 10 weeks. A) Male Mybpc3^-/-^ mice (N=5) had a significantly lower body weight compared to wild-type male mice (N=5; P<0.000001). Female Mybpc3^-/-^ mice (N=4) had a significantly lower body weight compared to female wild-type mice (N=4; P=0.000131). B) Male Mybpc3^-/-^ mice (N=5) had a significantly lower mean GFP (EFP) weight compared to wild-type male mice (N=5; P=0.000085). Female Mybpc3^-/-^ mice (N=4) had a significantly lower mean GFP (POAT) weight compared to wild - type mice (N=4). C) Male EFP weights were normalized to BW to calculate an EFP / BW ratio to serve as an estimation of body composition. Two-way ANOVA analysis showed both male (N=8) and female (N=8) Mybpc3^-/-^ mice had a significantly lower GFP / BW ratio compared to wild-type males (N=8) and females (N=7; F (1, 15) = 37.52; P<0.00001). D) A qualitative histology study was performed with H&E, MT, ORO, and PAS in heart tissue isolated from wild-type and Mybpc3^-/-^ male mice fed a HFD for 10 weeks. E) A qualitative histology study was performed with H&E, MT, ORO, and PAS in heart liver tissue isolated from wild-type and Mybpc3^-/-^ male mice fed a HFD for 10 weeks. **** = P<0.00001. All error bars represent SD.

### A high fat diet boosts *Mybpc3^-/-^* cardiac respiration

Experiments to challenge cardiomyocyte and whole-body metabolism were repeated in *Mybpc3**^−/−^*** males fed a HFD for 10 weeks to assess the impact of altered lipid stores due to exogenous LCFA supplementation. One-way ANCOVA analysis of O_2_ consumption in *Mybpc3**^−/−^*** males compared to wild-type males showed no significant effects (F = 0.9538, p = 0.3543) (Table 2). However, there was no glucose catabolism-driven spike in *Mybpc3**^−/−^*** RER during an acute exercise challenge, suggesting highly metabolic tissues were more reliant on FAO OXPHOS instead of glycolysis **(Figure 10A)**. Although *Mybpc3**^−/−^*** resting RER was significantly higher, wild-type mice experienced a glucose-driven spike in RER during an acute exercise challenge, as expected and seen in both genotypes on a regular diet **(Figure 6A)**. *Mybpc3**^−/−^*** males had significantly lower glucose levels, but higher ketone body levels, pre- and post-exercise challenge compared to wild-type mice, despite supplementation with exogenous LCFAs (Figure 10B & C).

**Figure 10.**
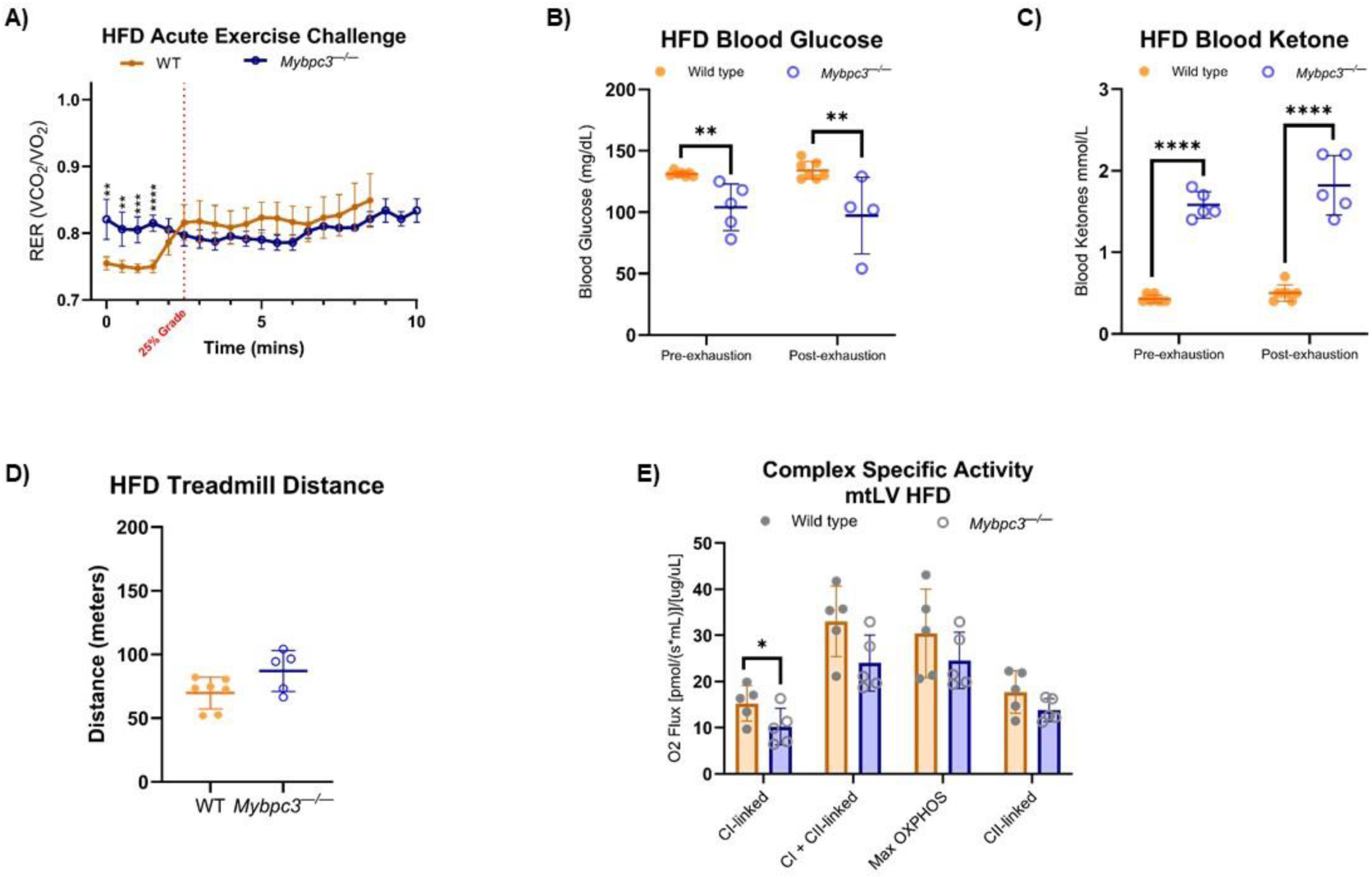
Wild-type and Mybpc3^-/-^ mice were fed a HFD for 10 weeks. A) Exercise naïve Mybpc3^-/-^ and wild-type mice were subject to an acute exercise challenge, reported as an RER moving average. A UC analysis was performed; wild-type mice (N=7) had a peak area value of 6.835±0.036, and Mybpc3^-/-^ mice (N=5) had a peak area value of 8.044±0.023. Mybpc3^-/-^ mice had a significantly higher RER at the beginning of the exercise challenge (Q=0.0011; Q=0.0011; Q=0.0002; Q<0.00001). B) Two-way ANOVA analysis indicated that Mybpc3^-/-^ mice (N=5) had significantly lower blood glucose levels compared to wild-type mice (N=7), both pre- and post-exercise challenge (F (1, 6) = 39.59; P=0.0008). C) Two-way ANOVA analysis indicated that Mybpc3^-/-^ mice (N=5) had significantly lower blood ketone levels compared to wild-type mice (N=7), both pre- and post-exercise challenge. D) There was no significant difference in mean distance run between Mybpc3^-/-^ vs wild-type mice (unpaired t-test; P = 0.0614). E) Complex-specific activity was measured in mtLV collected from Mybpc3^-/-^ and wild-type mouse hearts, represented as an oxygen flux. Mybpc3^-/-^ mice (N=5) had significantly lower CI-linked activity compared to control counterparts (N=5; P=0.0377). CI + CII-linked (P=0.0966), maximu m OXPHOS (P=0.2372), and CII-linked (P=0.2073) oxygen flux was not significantly different between Mybpc3^-/-^ vs wild-type mtLV. * = P<0.05; ** = P<0.005; *** = P<0.0005; **** = P<0.00001. All error bars represent SD.

**Table 2.** One-way ANCOVA analysis was performed; assuming that body mass was a covariate of O_2_ Consumption. Genotype had no significant effect on O_2_ consumption.

**HFD *Mybpc3*<sup>-/-</sup> vs Wild-Type ANCOVA Analysis**
|  | Sum of Squares | df | F | P-value |
| --- | --- | --- | --- | --- |
| **Genotype** | 8.95E+04 | 1.0 | 0.9538 | 0.3543 |
| **Body Mass** | 1.22E+05 | 1.0 | 1.3014 | 0.2834 |
| **Residual** | 8.45E+05 | 27.0 | NA | NA |

Complex-specific activity was measured in mitochondria isolated from cardiac tissue in animals fed a HFD. *Mybpc3**^−/−^*** mice had significantly reduced CI-linked respiration, but CI + CII-linked, maximum OXPHOS, and CII-linked respiration were not significantly different from wild-type respiration **(Figure 10E)**.

## Discussion

The findings presented in this study indicate that *Mybpc3**^−/−^*** mice, particularly males, exhibit a notable shift toward increased reliance on glucose catabolism for ATP production in both cardiac and skeletal muscle. *Mybpc3**^−/−^*** mice showed significantly decreased oxygen consumption and energy expenditure compared to wild-type mice. This is consistent with previous studies performed with human myocardium isolated from patients with cardiac hypertrophy, which also report shifts in energy metabolism away from FAO, toward glucose catabolism OXPHOS in cardiac tissue [35,36]. However, the observation that *Mybpc3**^−/−^*** mice display a reduction in whole-body expenditure warrants further investigation to determine the underlying mechanisms of this apparent systemic shift in bioenergetic preference for glycolysis. Although, metabolic phenotypes could be affected by activity levels, our treadmill exercise data **(Figure 6A)** showed that *Mybpc3**^−/−^*** mice run for the same duration as wild-type mice under an identical exercise protocol. Additionally, if the LVH phenotype in *Mybpc3**^−/−^*** mice were restricting pulmonary function and thus limiting exercise capacity, we would have expected to see significant differences in treadmill performance and time to exhaustion. Since *Mybpc3**^−/−^*** mice performed similarly to wild-type mice, we can infer that the altered metabolic phenotype is not primarily driven by decreased exercise tolerance caused by LVH [1]. This suggests that the RER metabolic expenditure differences observed in *Mybpc3**^−/−^*** mice are not due to differences in physical activity levels, but rather, an inherent systemic metabolic shift due to cardiac hypertrophy.

We believe that the presence of LVH in *Mybpc3**^−/−^*** mice caused the metabolic reprogramming observed in other highly metabolic tissues. Our histology and functional mitochondrial studies showed similar shifts in metabolic processes. Although it is possible that reduced oxygen availability may contribute to this metabolic shift, our data also suggest that supplementation of exogenous FFAs reversed this phenotype in myocardium **(Figure 10E)**, indicating that lipid utilization and storage could be a key modulator of systemic metabolism in LVH. Previous studies have also provided evidence that increasing the levels of carnitine palmitoyl transporters (CPTs) as a therapeutic target increases fatty acid uptake by the mitochondria, thereby increasing LCFA oxidation and reducing cardiac lipotoxicity in patients with various causes of LVH [9,13,37–39]. Another proposed mitochondrial target, α-ketoglutarate dehydrogenase (KGDH), synthesizes succinyl-CoA from α-ketoglutarate as one of the NADH-generating steps of the tricarboxylic acid (TCA) cycle [40–42]. Since this reaction is NAD^+^ dependent, KGDH acts as a mitochondrial redox sensor to reversibly suppress mitochondrial activity under extreme oxidative stress [41]. More recently, KGDH has been shown to play a role in facilitating H_2_O_2_ elimination in response to CI blockades in the heart [40]. Therefore, it is possible for cardiomyocytes to generate ATP with OXPHOS, with the right substrates. KGDH expression levels, stability, and activity studies are necessary to determine whether exogenous supplementation of LCFAs counteracts mitochondrial suppression via this mechanism.

This is corroborated by our *in vitro* O_2_ consumption data **(Figure 7A)**, which indicate that the altered metabolic state in *Mybpc3**^−/−^*** mtLV is associated with diminished oxidative capacity, further supporting a shift away from OXPHOS and towards glycolysis **(Figure 7B)**. Interestingly, though, the provision of exogenous LCFA both *in vitro* and *in vivo* reversed many metabolic abnormalities observed in *Mybpc3**^−/−^*** mice, suggesting that their systemic metabolic dysfunction, may, in part, be due to impaired lipid utilization outside cardiac tissue. A HFD resulted in loss of hepatic and cardiac lipotoxicity **(Figure 9D & 9E)** with increased mitochondrial OXPHOS capacity **(Figure 10E)**, but did not fully rescue the *Mybpc3**^−/−^*** systemic phenotypes of increased baseline RER **(Figure 10A)**, altered body composition **(Figure 9C)**, nor blood glucose and ketone levels indicative of metabolic duress **(Figure 10B & 10C)**. It is possible that LCFAs supplemented by a HFD may not be utilized by the heart, but rather support FAO in other highly metabolic tissues, such as skeletal muscle. Therefore, the heart has preferential use of most, if not all, bioavailable glucose, while other tissues have access to LCFAs, such that the bioavailability of *usable* OXPHOS precursors supersedes metabolic reprogramming in response to oxygen availability. The specific role of fatty acid, specifically LCFA, uptake and catabolism to acetyl-CoA in the progression of cardiomyopathies and heart failure warrants further investigation *in vivo* to track fatty acid vs glucose uptake in heart, liver, skeletal muscle, brain, and kidney. We suspect that a study like this would show that *Mybpc3^−/−^* liver is forced to mostly use FAO as its fuel source. Not only are ketone levels constitutively high in *Mybpc3^−/−^* mice – whether glycolytically challenged during a fasting **(Figure 5B)** or during acute exercise **(Figure 6C)** – but RER data indicated higher reliance on glucose in muscle, heart, kidney, liver, and brain tissues. The only way for the body to produce ketones is through liver FAO [43], supporting our hypothesis that compromised heart muscle “forces” a systemic metabolic phenotype; directly affecting liver FAO while indirectly affecting skeletal muscle FAO.

Our study using an established mouse model of LVH provides clinically relevant insights into the metabolic consequences of inherited and acquired forms of CVD that result in cardiac hypertrophy. Our observation of systemic metabolic disturbances in *Mybpc3^−/−^* mice due to cardiac hypertrophy may help identify new biomarkers or therapeutic targets that address not only the heart, but also the systemic metabolic consequences of cardiac dysfunction. The use of serum acylcarnitine profiling in this study provided insight into the fatty acids primarily not used for heart FAO, then skeletal muscle FAO **(Figure 5C – 5H)** [29]. Acylcarnitines are formed inside the mitochondria, then shipped out of the cell when not used as a precursor for acetyl-CoA synthesis via FAO. Since we found a significant increase in several long chain-acylcarnitines in *Mybpc3^−/−^* serum, we can surmise that there is an FAO bottleneck of some sort in heart and/or skeletal muscle.

These profiles are commonly used to screen for inborn errors of metabolism (IEMs) in humans, providing a comprehensive profile of FAO disorders, which cause systemic metabolic dysfunction [44]. Patients with IEMs, particularly those with FAO disorders or hypercholesterolemia, are frequently monitored for the development of CVD [45,46]. Interestingly, we found that the acyl-carnitine profiles of *Mybpc3^−/−^* mice closely match those observed in patients with Short-chain Acyl-CoA Dehydrogenase Deficiency (SCAHD), a type of IEM characterized by significant increases in C4OH. This similarity supports the possibility that inherited genetic variants that cause hypertrophy in the heart may disrupt systemic metabolism in a manner analogous to IEMs, which have clinical implications in liver, heart, kidneys, brain, and skeletal muscle [46]. This observation is significant, as it suggests that mutations restricted to cardiac tissue can have wide-ranging effects on metabolic processes throughout the entire organism.

This study provides evidence that LVH caused by loss of the cardiac-specific gene, *Mybpc3*, can indeed induce systemic metabolic dysregulation, extending beyond the heart and affecting whole-body energy balance. This study also provides a valuable framework for studying the systemic metabolic alterations that accompany inherited forms of LVH. The findings underscore the importance of considering the whole-body metabolic effects of cardiac mutations thought to only affect the heart, and suggest that therapeutic strategies targeting broader metabolic pathways may be beneficial in managing patients with LVH.

## Supporting information

Supplemental Data S1-S5 and Acylcarnitine Table

## Notes

### Competing Interest Statement

The authors have declared no competing interest.

### Funding

- National Heart Lung and Blood Institute: R01 HL173989-02, R01 HL153236-04, R01 HL175803

