## Supplemental Data S1-S5 and Acylcarnitine Table for "Systemic Metabolic Rewiring in a Mouse Model of Left Ventricular Hypertrophy"

### *Supplementary Material*

#### 1 Supplementary Data

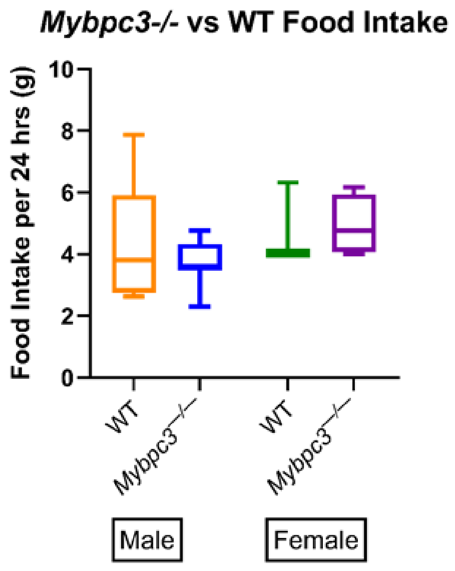


**Supplementary Figure 1.** Food intake data are represented as a 24 hr average, measured over the course of 24 hrs. There was no significant difference between male *Mybpc3*$^{-/-}$ (N=7) and wild-type males (N=6; P=0.4515). There was no significant difference between female *Mybpc3*$^{-/-}$ mice (N=4) and wild-type mice (N=3; P=0.8731). Error bars represent SD.

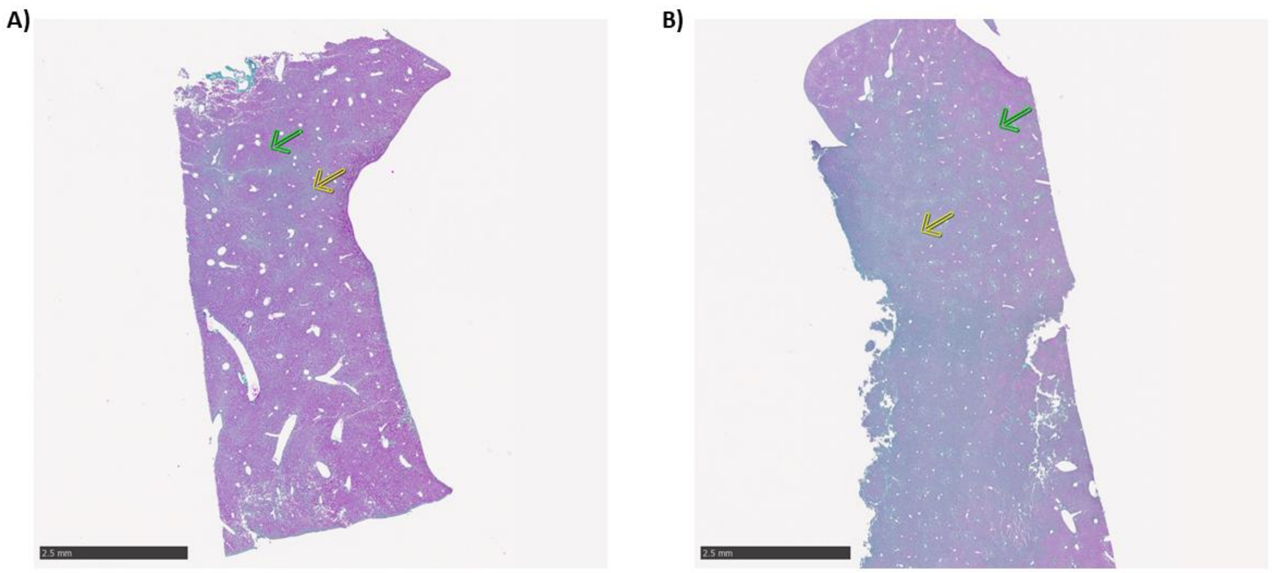


**Supplemental Figure 2. A)** whole-slide scan of liver stained with PAS isolated from a wild-type mouse. The green arrow indicates an example of a glycogen-rich region in the liver. The yellow arrow indicates a glycogen-deplete region in the liver. **B)** Whole-slide scan of liver section stained with PAS isolated from a *Mybpc3*$^{-/-}$ mouse. The green arrow indicates an example of a glycogen-rich region in the liver. The yellow arrow indicates a glycogen-deplete region in the liver.

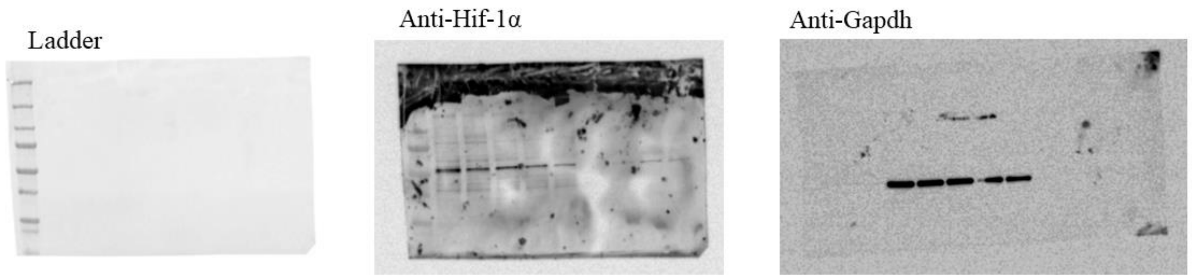


**Supplemental Figure 3.** Full membrane images for anti-Hif-1α, anti-Gapdh, and ladder immunoblots.

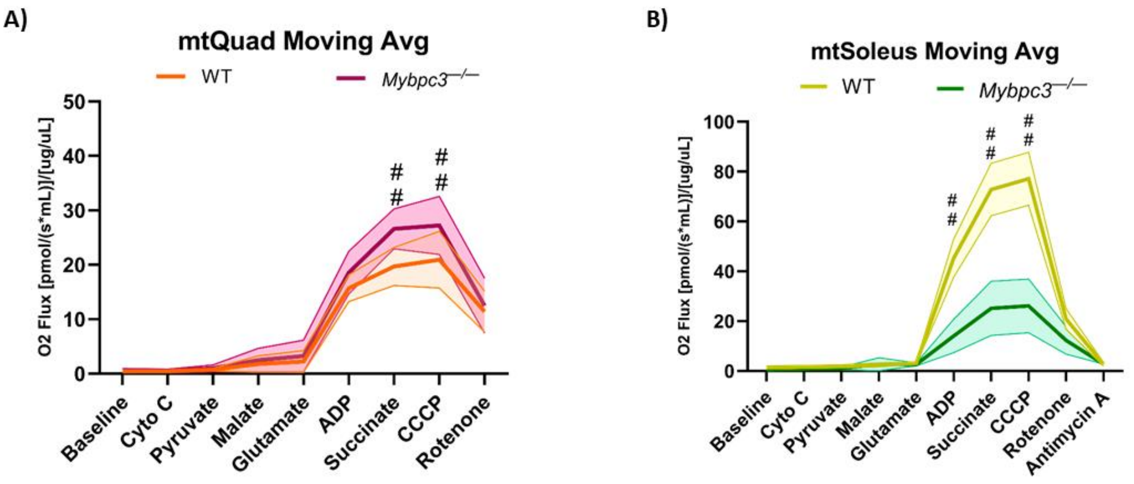


**Supplemental Figure 4. A)** Oxygen consumption of mtQuad in response to various substrates was plotted as a moving mean. Multiple t-test analysis showed *Mybpc3*$^{-/-}$ mtQuad (N=6) had significantly higher oxygen consumption with treatment of succinate (Q=0.001188) and CCCP (Q=.0001890) compared to wild-type mtQuad (N=6). **B)** Oxygen consumption of mtSoleus in response to various substrates was plotted as a moving mean. Multiple t-test analysis showed *Mybpc3*$^{-/-}$ mtSoleus (N=6) had significantly higher oxygen consumption with treatment of ADP (Q=0.000288), Succinate (Q=0.000288), and CCCP (Q=0.000288) compared to wild-type mice (N=6). Error bars (shaded region) represents SD; ## = Q<0.001.

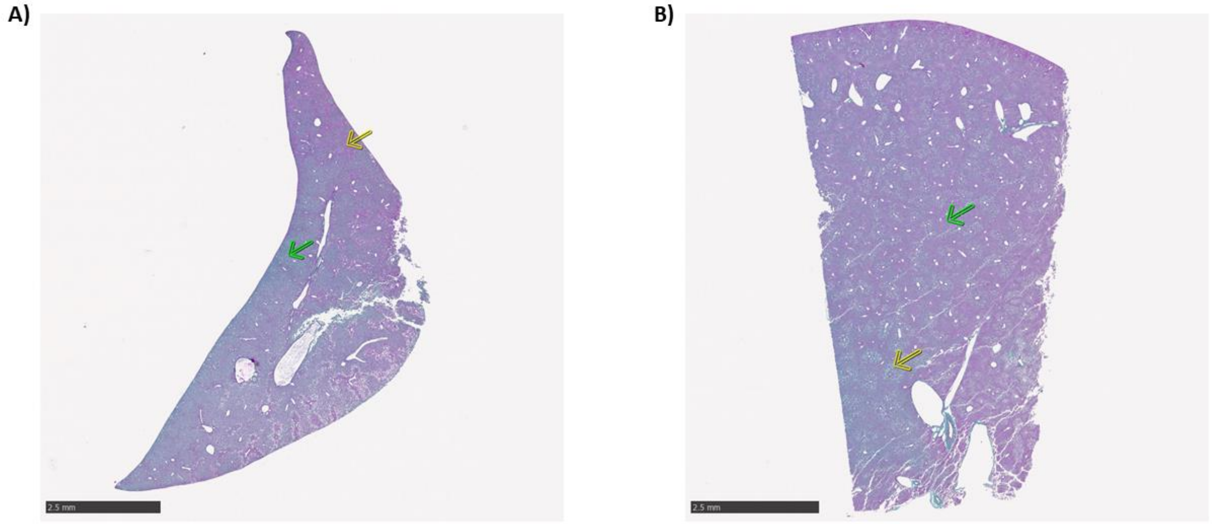


**Supplemental Figure 5.** *Mybpc3*$^{-/-}$ and wild-type mice were fed a HFD for 10 weeks. **A)** whole-slide scan of liver stained with PAS isolated from a wild-type mouse. The green arrow indicates an example of a glycogen-rich region in the liver. The yellow arrow indicates a glycogen-deplete region in the liver. **B)** Whole-slide scan of liver section stained with PAS isolated from a *Mybpc3*$^{-/-}$ mouse. The green arrow indicates an example of a glycogen-rich region in the liver. The yellow arrow indicates a glycogen-deplete region in the liver.

### 2 Supplementary Tables and Figures

#### 2.1 Circulating Acylcarnitine Concentrations

***Mybpc3-/-*** **vs Wild-type Acylcarnitine Profiling**

| | Wild-type | | | | Mybpc3-/- | | | | | |
|---|---|---|---|---|---|---|---|---|---|---|
| **Substrate Name** | **Final Concentration** | | | **WT Avg** | **Final Concentration** | | | **Mybpc3-/- Avg** | **P-value** | **FC (Mybpc3-/- / WT)** |
| C0 | 19.7816 | 25.7394 | 18.5903 | 21.3704 | 24.2841 | 16.8109 | 19.4745 | 20.1898 | 0.7235 | 0.94 |
| C2 | 18.7453 | 30.0778 | 21.6525 | 23.4919 | 25.6996 | 22.2359 | 22.7533 | 23.5629 | 0.9850 | 1.00 |
| C3 | 0.7393 | 1.0653 | 0.8966 | 0.9004 | 0.9667 | 0.7619 | 0.8212 | 0.8499 | 0.6758 | 0.94 |
| C4 | 0.4153 | 0.4718 | 0.3996 | 0.4289 | 0.6492 | 0.3348 | 0.3526 | 0.4455 | 0.8810 | 1.04 |
| C4OH | 0.1561 | 0.1467 | 0.1044 | 0.1357 | 0.2764 | 0.2225 | 0.2282 | 0.2424 | 0.0103 | 1.79 |
| C5 | 0.1270 | 0.2717 | 0.2116 | 0.2034 | 0.3232 | 0.2011 | 0.1939 | 0.2394 | 0.5772 | 1.18 |
| C5_isomers | 0.0893 | 0.1449 | 0.0935 | 0.1092 | 0.1024 | 0.0738 | 0.0762 | 0.0841 | 0.2795 | 0.77 |
| C5:1 | 0.0048 | 0.0074 | 0.0064 | 0.0062 | 0.0125 | 0.0081 | 0.0070 | 0.0092 | 0.1789 | 1.48 |
| C5OH | 0.0000 | 0.0717 | 0.0568 | 0.0428 | 0.0851 | 0.0592 | 0.0612 | 0.0685 | 0.3338 | 1.60 |
| C6 | 0.0596 | 0.1175 | 0.0902 | 0.0891 | 0.1108 | 0.1100 | 0.0890 | 0.1033 | 0.4794 | 1.16 |
| C6OH | 0.0081 | 0.0095 | 0.0078 | 0.0085 | 0.0107 | 0.0104 | 0.0110 | 0.0107 | 0.0155 | 1.26 |
| C8 | 0.0211 | 0.0161 | 0.0105 | 0.0159 | 0.0130 | 0.0129 | 0.0072 | 0.0110 | 0.2491 | 0.69 |
| C8:1 | 0.0086 | 0.0083 | 0.0059 | 0.0076 | 0.0042 | 0.0045 | 0.0026 | 0.0038 | 0.0210 | 0.50 |
| C8OH | 0.0133 | 0.0143 | 0.0120 | 0.0132 | 0.0145 | 0.0129 | 0.0116 | 0.0130 | 0.8609 | 0.98 |
| C10 | 0.0000 | 0.0000 | 0.0000 | 0.0000 | 0.0000 | 0.0000 | 0.0000 | 0.0000 | NA | NA |
| C10:1 | 0.0000 | 0.0000 | 0.0000 | 0.0000 | 0.0000 | 0.0000 | 0.0000 | 0.0000 | NA | NA |
| C10:2 | 0.0000 | 0.0000 | 0.0000 | 0.0000 | 0.0000 | 0.0000 | 0.0000 | 0.0000 | NA | NA |
| C10OH | 0.0087 | 0.0102 | 0.0063 | 0.0084 | 0.0058 | 0.0061 | 0.0037 | 0.0052 | 0.0788 | 0.62 |
| C12 | 0.0127 | 0.0203 | 0.0075 | 0.0135 | 0.0120 | 0.0070 | 0.0034 | 0.0075 | 0.2489 | 0.55 |
| C12:1 | 0.0118 | 0.0142 | 0.0095 | 0.0118 | 0.0107 | 0.0127 | 0.0063 | 0.0099 | 0.4528 | 0.84 |
| C12OH | 0.0000 | 0.0000 | 0.0000 | 0.0000 | 0.0000 | 0.0000 | 0.0000 | 0.0000 | NA | NA |
| C14 | 0.0411 | 0.1010 | 0.0646 | 0.0689 | 0.0711 | 0.0620 | 0.0668 | 0.0666 | 0.9039 | 0.97 |
| C14:1 | 0.0737 | 0.0929 | 0.0524 | 0.0730 | 0.1095 | 0.0763 | 0.0646 | 0.0835 | 0.5886 | 1.14 |
| C14:1OH | 0.0000 | 0.0000 | 0.0000 | 0.0000 | 0.0000 | 0.0000 | 0.0000 | 0.0000 | NA | NA |
| C14:2 | 0.0262 | 0.0392 | 0.0242 | 0.0299 | 0.0308 | 0.0237 | 0.0171 | 0.0239 | 0.3841 | 0.80 |
| C14OH | 0.0000 | 0.0000 | 0.0000 | 0.0000 | 0.0000 | 0.0000 | 0.0000 | 0.0000 | NA | NA |
| C16 | 0.0401 | 0.1127 | 0.0747 | 0.0758 | 0.1138 | 0.1063 | 0.1342 | 0.1181 | 0.1343 | 1.56 |
| C16:1 | 0.0266 | 0.0524 | 0.0301 | 0.0364 | 0.0693 | 0.0569 | 0.0419 | 0.0560 | 0.1572 | 1.54 |
| C16:1OH | 0.0033 | 0.0050 | 0.0038 | 0.0040 | 0.0064 | 0.0060 | 0.0067 | 0.0064 | 0.0127 | 1.58 |
| C16OH | 0.0069 | 0.0134 | 0.0093 | 0.0099 | 0.0161 | 0.0125 | 0.0147 | 0.0144 | 0.1029 | 1.46 |
| C18 | 0.0738 | 0.0531 | 0.0247 | 0.0505 | 0.0864 | 0.0503 | 0.1080 | 0.0816 | 0.2319 | 1.61 |
| C18:1 | 0.0173 | 0.0468 | 0.0219 | 0.0287 | 0.1014 | 0.0634 | 0.0808 | 0.0819 | 0.0205 | 2.86 |
| C18:1OH | 0.0418 | 0.0570 | 0.0419 | 0.0469 | 0.0389 | 0.0477 | 0.0469 | 0.0445 | 0.6992 | 0.95 |
| C18:2 | 0.0819 | 0.0955 | 0.0729 | 0.0834 | 0.1380 | 0.0849 | 0.1180 | 0.1136 | 0.1470 | 1.36 |
| C18:2OH | 0.0235 | 0.0418 | 0.0277 | 0.0310 | 0.0465 | 0.0320 | 0.0357 | 0.0381 | 0.3722 | 1.23 |
| C18OH | 0.0000 | 0.0000 | 0.0000 | 0.0000 | 0.0000 | 0.0000 | 0.0000 | 0.0000 | NA | NA |
